# Range-wide spatial mapping reveals convergent character displacement of bird song

**DOI:** 10.1101/560276

**Authors:** Alexander N. G. Kirschel, Nathalie Seddon, Joseph A. Tobias

**Affiliations:** University of Cyprus, Department of Biological Sciences, PO Box 20537, Nicosia 1678, Cyprus.; Edward Grey Institute, Department of Zoology, University of Oxford, South Parks Road, Oxford OX1 3PS, UK.; Department of Life Sciences, Imperial College London, Silwood Park, Buckhurst Road, Ascot, Berkshire, SL5 7PY, UK.

**Keywords:** animal communication, Amazonia, interspecific competition, phenotypic evolution, signal convergence, social selection

## Abstract

A long-held view in evolutionary biology is that character displacement generates divergent phenotypes in closely related coexisting species to avoid the costs of hybridisation or ecological competition, whereas an alternative possibility is that signals of dominance or aggression may instead converge to facilitate coexistence among ecological competitors. Although this counter-intuitive process—termed convergent agonistic character displacement—is supported by recent theoretical and empirical studies, the extent to which it drives spatial patterns of trait evolution at continental scales remains unclear. By modeling variation in song structure of two ecologically similar species of *Hypocnemis* antbird across western Amazonia, we show that their territorial signals converge such that trait similarity peaks in the sympatric zone, where intense interspecific territoriality between these taxa has previously been demonstrated. We also use remote sensing data to show that signal convergence is not explained by environmental gradients and is thus unlikely to evolve by sensory drive (i.e. acoustic adaptation to the sound transmission properties of habitats). Our results suggest that agonistic character displacement driven by interspecific competition can generate spatial patterns opposite to those predicted by classic character displacement theory, and highlight the potential role of social selection in shaping geographical variation in signal phenotypes of ecological competitors.

## Introduction

Classical evolutionary theory suggests that mating signals of closely related species diverge when populations interact in geographical contact (sympatry) to reduce the costs of maladaptive hybridisation or reproductive interference (Dobzhansky 1951, Butlin 1987, Servedio and Noor 2003, Groning and Hochkirch 2008, Pfennig and Pfennig 2009). This concept, widely referred to as reproductive character displacement (RCD) (Butlin 1987, Gerhardt 1994, Pfennig and Pfennig 2009), is thought to explain patterns of divergence in both signals and signal recognition systems, particularly in anurans (Gerhardt 1994, Pfennig and Pfennig 2005, Lemmon 2009), and birds (Saetre et al. 1997, Kirschel et al. 2009). An alternative hypothesis, first suggested by West-Eberhard (West-Eberhard 1983), is that agonistic signals functioning in social competition may also diverge to reduce the costs of misdirected interspecific aggression—a process now termed agonistic character displacement (ACD) (Grether et al. 2009, Grether et al. 2013, Grether et al. 2017). Standard forms of both RCD and ACD predict signal divergence in sympatry, making them difficult to disentangle through observational studies (Kirschel et al. 2009, Tobias et al. 2011, Tobias et al. 2014) and thus their relative contribution to patterns of trait evolution is often unclear (Grether et al. 2009, Grether et al. 2013, Grether et al. 2017).

RCD is by definition exclusively divergent, whereas a key discriminating feature of ACD is that, under certain conditions, social signals of competing species may theoretically converge to mediate interspecific territoriality and facilitate competitor recognition (Cody 1969, Grether et al. 2009, Tobias and Seddon 2009, Laiolo 2012, Grether et al. 2017). Evidence for convergent ACD has been accumulating over recent years. For example, theoretical studies have concluded that socially mediated signal convergence among ecological competitors is plausible (Scheffer and van Nes 2006, Grether et al. 2009), while empirical studies have reported community-wide or interspecific patterns consistent with this hypothesis (Laiolo 2012, Tobias et al. 2014, Tobias et al. 2014, Leighton et al. 2018). Nonetheless, demonstrating that species interactions drive convergence in social signals remains challenging because similar effects can be produced by (i) hybridization, (ii) heterospecific copying of learned signals, and (iii) ecological adaptation to shared habitats (Slabbekoorn and Smith 2002, Seddon 2005, Kirschel et al. 2009, Kirschel et al. 2009, Tobias et al. 2010), all of which can increase signal similarity between sympatric taxa (Tobias et al. 2010). One way of accounting for these factors is to assess whether convergence can reverse the classic spatial pattern associated with character displacement—i.e. a gradient of increasing trait similarity from allopatry to sympatry (Servedio 2004)—while controlling for underlying environmental variation (Goldberg and Lande 2006). However, few studies have adopted this approach, partly because it is difficult to map variation of signal traits in relation to habitat at continental scales (Smith et al. 2013).

To address this issue, we analysed the acoustic structure of song in two species of antbird—Peruvian warbling-antbird (*Hypocnemis peruviana*) and yellow-breasted warbling-antbird (*H. subflava*)—which appear to use convergent songs to defend interspecific territories where their ranges overlap in Western Amazonia (Tobias and Seddon 2009, Tobias et al. 2011). Despite their similar songs, these non-sister taxa are separated by 6.8% sequence divergence in mitochondrial DNA (Tobias et al. 2008), and experiments in captivity have demonstrated that females can discriminate between individuals and species with high accuracy on the basis of song (Seddon and Tobias 2010). Accordingly, they rarely if ever hybridise (Tobias and Seddon 2009). Moreover, *Hypocnemis* antbirds are tracheophone suboscine passerines, a group for which a range of observational and experimental evidence suggests that songs are genetically determined (innate) (Seddon and Tobias 2006, Tobias et al. 2010, Tobias et al. 2012, Tobias et al. 2014, Touchton et al. 2014). Accordingly, the advantage of the *Hypocnemis* system is that song convergence is unlikely to be caused by either hybridization or heterospecific copying, two factors which may explain similar patterns of variation reported in oscine passerines (songbirds with learned songs) (Grether et al. 2009, Laiolo 2012, Reif et al. 2015).

Previous work has shown that *H. peruviana* responds more aggressively to playback of *H. subflava* territorial song in sympatry than allopatry (Tobias and Seddon 2009). However, the extent to which this behaviour reflects spatial convergence in song was not clear because the findings were based on observations from just three sites, one in sympatry and one for each species in allopatry, reporting no consistent pattern of divergence or convergence in acoustic traits (Tobias and Seddon 2009). In addition, previous comparisons between sympatry and allopatry did not take into account the role of environmental factors such as vegetation density, which can influence song evolution by acoustic adaptation to the sound transmission properties of the signaling environment (Seddon 2005, Kirschel et al. 2009, Tobias et al. 2010).

We conducted field surveys and compiled sound files from public and private archives to estimate song variation over the entire range of *H. peruviana* and *H. subflava*, sampling widely within both sympatric and allopatric zones. This approach included all known subspecies, some of which are allopatric and others partially sympatric. We first compared songs of the different subspecies to assess whether patterns of signal variation were predicted by geographical contact between taxa. We then focused on the taxon with the widest range and deepest sampling (*H. peruviana*) to assess whether its song converges from allopatry towards the contact zone, where it interacts with *H. subflava*. Finally, to control for patterns of convergence attributable to acoustic adaptation, we incorporated environmental data from remote sensing into spatial models of signal variation.

## Material and methods

### (a) Study system

All *Hypocnemis* antbirds inhabit dense forest in the Amazon basin and foothills of the Andes (Isler et al. 2007, Tobias et al. 2008). The ranges of *H. peruviana* and *H. subflava* overlap in Southwest Peru, West Brazil, and North Bolivia, with the region of sympatry spanning □1100 km at its widest point and covering over 150,000 km^2^ (Tobias and Seddon 2009). They sing at similar heights above ground, and occupy similar foraging niches, including near-identical foraging techniques, diet, and biometric traits (Tobias et al. 2008, Tobias and Seddon 2009). Both species compete for territories in ecotonal or successional vegetation, although otherwise they are partially segregated into two core habitats—*H. peruviana* mainly in *terra firme* forest; *H. subflava* mainly in *Guadua* bamboo (Tobias and Seddon 2009)—which are distributed in a complex mosaic throughout the contact zone (Smith and Nelson 2011).

*H. peruviana* and *H. subflava* are highly divergent in terms of plumage signals (Tobias and Seddon 2009), but some of their acoustic signals are very similar (figure 1). In particular, both species use a multi-note sex-specific song that mediates mate-choice and territorial defence in both sexes and is given either as solos or male-led duets (Seddon and Tobias 2006, Tobias and Seddon 2009). We focus on the male song because females sing less frequently, and often overlapping with male songs (Tobias and Seddon 2009), making it difficult to collate a large sample of unmasked high-quality sound files for analysis.

**Figure 1.**
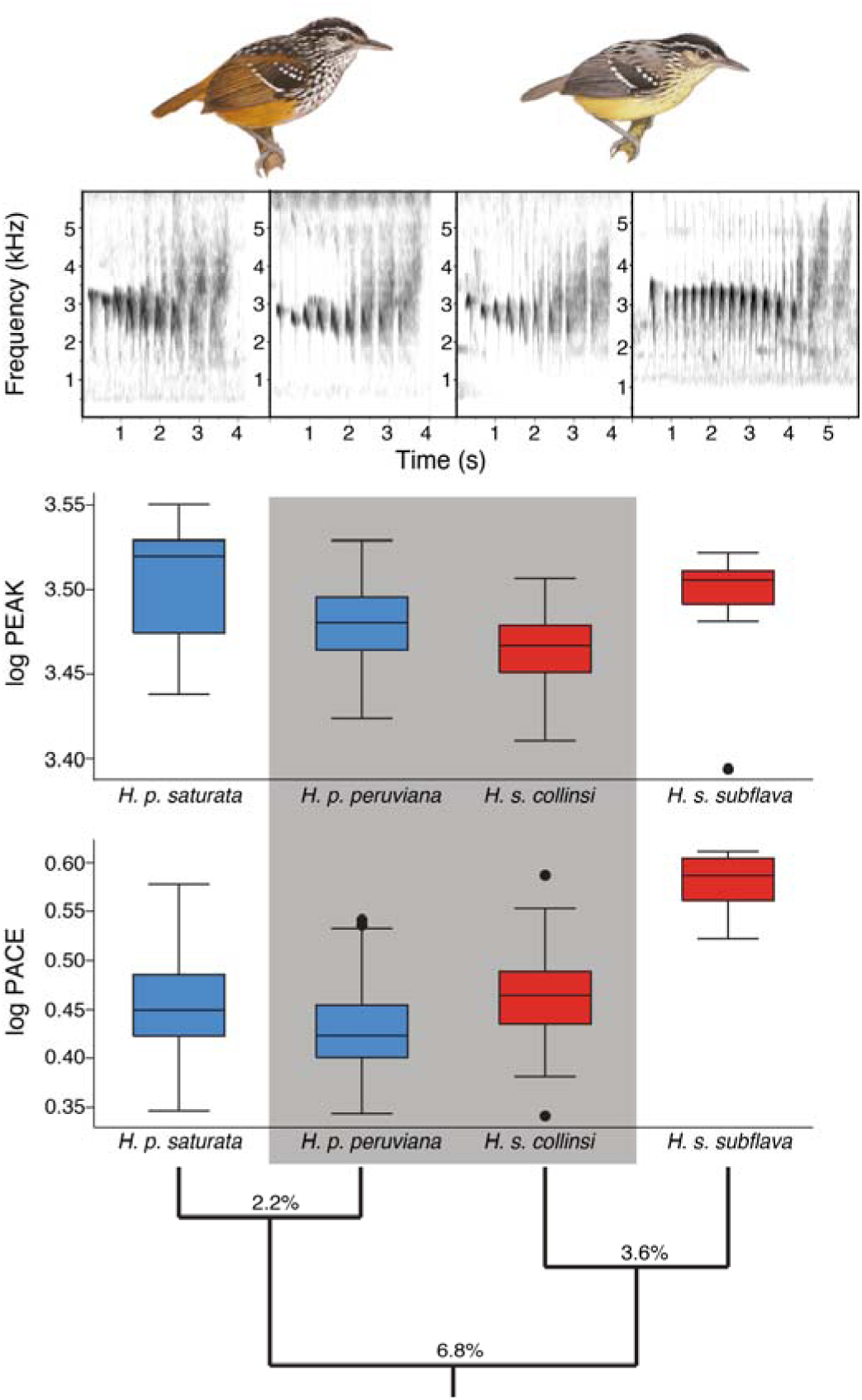
Songs of sympatric *H. p. peruviana* and *H. s. collinsi* (grey) are more similar in peak frequency and pace than predicted from their genetic distance (% mtDNA sequence divergence). Songs of *H. s. collinsi* are broadly similar to those of *H. p. peruviana* (sympatric), but differ significantly in both peak frequency and pace from conspecific *H. s. subflava* (allopatric). Levels of significance following FDR correction: * P < 0.05, ** P < 0.01, *** P < 0.001, ns = not significant. Spectrograms show representative songs of (left to right) *H. p. saturata, H. p. peruviana, H. s. collinsi*, and *H. s. subflava.* Songs of *peruviana* and *collinsi* were both recorded in the contact zone. Illustrations are reproduced with the permission of Lynx Edicions (del Hoyo 2018).

### (b) Song sampling and analysis

We obtained song recordings of all four subspecies (*H. p. peruviana, H. p. saturata, H. s. subflava, H. s. collinsi*) from a total of 86 different localities distributed throughout their geographical ranges (30 sympatric sites; 58 allopatric sites). We recorded 193 individuals during fieldwork in Brazil, Bolivia, Ecuador and Peru between 2001 and 2011. Field recordings were made using a Marantz PMD 661 or Sound Devices 722 portable recorder with a Sennheiser microphone (MKH 8050 or ME67-K3U). Additional recordings were compiled from global sound archives, including xeno-canto (http://xeno-canto.org; 2 recordings) and Macaulay Library (http://macaulaylibrary.org; 18 recordings), as well as private collections (141 recordings; see electronic supplementary material for a discussion of sampling methods and limitations, and a complete list of recording localities [table S1]). All files were saved as 44.1 kHz wav files.

We measured song structure using RAVEN PRO 1.4 (Cornell Lab of Ornithology, Ithaca, NY), sampling at least six high quality songs per individual where possible. The final dataset contained ∼40,000 measurements from 1661 songs of 355 individuals, with a mean (± SD) of 4.8 ± 2.2 songs sampled per individual, and 4.1 ± 5.5 individuals sampled per site. We processed these songs using the MatLab signal processing toolbox (Mathworks, Natick, MA), automatically extracting a total of 22 spectral and temporal acoustic measures for analysis (table S3; see electronic supplementary material for a full description of song analyses). In addition to analyses performed using all 22 acoustic measures, we followed previous studies of species with similar songs (Kirschel et al. 2009, Tobias et al. 2014, Nwankwo et al. 2018) by performing separate analyses on two of those measures, mean note peak frequency (hereafter, peak frequency) and overall song pace (hereafter, pace). Peak frequency was generated by calculating peak frequencies for each note of the song, and then taking the average across the entire song. Pace was calculated by dividing the number of notes minus 1, by the song duration minus mean note duration (see (Kirschel et al. 2009), table S3).

### (c) Quantifying environmental gradients

Previous studies based on a Geographic Information Systems (GIS) approach have shown that remotely sensed environmental data can predict both Amazonian tree species composition and song structure in rainforest understorey birds (Saatchi et al. 2008, Kirschel et al. 2009, Kirschel et al. 2011). This makes sense because GIS vegetation indices relate to canopy structure, which is known to regulate illumination, and thus vegetation growth and leaf density, in the understorey (Jennings et al. 1999). In accordance with the sensory drive hypothesis, the vegetation structure and density of the understorey has been shown to affect signal transmission in Amazonian birds (Tobias et al. 2010).

We used an established set of GIS methods (Saatchi et al. 2008, Kirschel et al. 2009, Kirschel et al. 2011) to quantify environmental variation across the geographic ranges of our study taxa. Specifically, we estimated elevation using Global Multi-resolution Terrain Elevation Data 2010 (https://topotools.cr.usgs.gov/gmted_viewer/), at a spatial resolution of 30 arc-seconds (Root Mean Square Error range: 24–42 m), and estimated climate using bioclimatic variables (annual mean temperature [BIO01] and annual precipitation [BIO12]) from the WorldClim Database (Hijmans et al. 2005). Both elevation and climate data were obtained at a 1-km^2^ resolution. To estimate habitat structure, we extracted two variables remotely sensing from the Moderate Resolution Imaging Spectroradiometer (MODIS; http://modis.gsfc.nasa.gov) at a resolution of 250 m: (i) Vegetation Continuous Field (VCF) (Hansen et al. 2002) for 2010 and (ii) Enhanced Vegetation Index (EVI) collected in August-September 2011. These time periods were chosen because of their relevance to the time of greatest fieldwork effort (see electronic supplementary material for further details and vegetation rasters [figure S1, S2]). The VCF product (https://modis.gsfc.nasa.gov/data/dataprod/mod44.php) represents percent tree cover, a relevant environmental variable in terms of signal transmission because it is theoretically related to habitat density, and has previously been shown to predict variation in song pitch of tropical forest birds (Kirschel et al. 2009, Kirschel et al. 2011). EVI is a composite property of leaf area, chlorophyll and canopy structure, considered more sensitive, at least in high biomass regions such as rainforest, to variation in canopy structure (Huete et al. 2002), which is typically saturated in MODIS’ other vegetation index, the normalized difference vegetation index (NDVI) (Phillips et al. 2008). To address cloud contamination—a problem with most remotely sensed data (Thomassen et al. 2010)—we substituted in the mean 5×5 grid values for 125 points for the EVI and VCF measures.

As we are interested in song convergence towards the contact zone, and the extent to which isolation-by-distance (Wright 1943) explains patterns of variation in allopatry, we correlated song variation to a linear measure of distance from the contact zone. This linear measurement is appropriate for our study because of the reasonably continuous nature of Amazonian rainforest within the global distribution of *H. peruviana*. We predicted that songs would show a gradual pattern of divergence or convergence from the zone of sympatry because of the effects of gene flow and isolation by distance (Tobias and Seddon 2009). We calculated the distance of *H. peruviana* recording localities from the closest edge of the contact zone using the near table function in ArcGIS 10.1 (Environmental Systems Research Institute 2011, Redlands, CA); all sympatric sites were given a zero value.

### (d) Spatial mapping of song structure

We plotted GPS coordinates of all recording sites, and categorized each site as either allopatric or sympatric with reference to published maps of the contact zone (Isler et al. 2007, Tobias and Seddon 2009). We then mapped the structure of *H. peruviana* song in relation to the structure of *H. subflava* songs in the contact zone. Specifically, we used the Inverse Distance Weighting function in ArcGIS 10.1 to map values for peak frequency and pace across the entire range of *H. peruviana* in western Amazonia (see electronic supplementary material for further details).

### (e) Statistical analyses

We conducted a principal components analysis (PCA) with varimax rotation on the correlation matrices of individual mean values (log-transformed) of the 22 acoustic traits extracted from song. This reduced the dimensionality of the song dataset and allowed us to quantify how the overall structure of songs varied within and between species and subspecies. The PCA generated seven principal components (PCs) with eigenvalues >1, which accounted for 85% of the variance in the song data. For factor loadings, see table S4.

We compared song differences between the four subspecies using analysis of variance (ANOVA) and Tukey’s HSD post hoc tests on PC scores extracted from song features using STATA 11 (StataCorp 2009). Our aim was to test whether *Hypocnemis* subspecies had more similar songs when they overlapped in geographical range (*H. p. peruviana* versus *H. s. collinsi*) in comparison with non-overlapping ranges (*H. p. saturata* versus *H. s. subflava*). Although the geographical ranges of *H. p. peruviana* and *H. s. collinsi* are only partially overlapping, we are focusing on innate acoustic signaling traits, and thus we expect that unimpeded gene flow from sympatry to allopatry obscures the classic stepped pattern of variation associated with character displacement (Tobias and Seddon 2009).

To examine this effect, and to determine the effect of sympatry with *H. subflava* on the song of *H. peruviana*, we ran generalized linear mixed models (GLMM) with Gaussian distribution and identity link function implemented in the lme4 (Bates et al. 2015) R Package. In these models, the dependent variable was one of nine aspects of song structure (averaged for each of the 198 individuals of *H. peruviana* included in the analysis; table S3). To control for environmental gradients, we included five geographical, climatic and habitat variables (see above) along with distance to contact zone as fixed factors in the GLMM. We also included recording site as a random effect to account for variation in sampling across sites. To account for multiple comparisons on seven PCs and further analyses of peak frequency and pace, we calculated false discovery rates based on those nine comparisons (Benjamini and Hochberg 1995, Verhoeven et al. 2005). Best models were selected based on the lowest corrected Akaike Information Criterion (AICc) score.

## Results

The overall taxonomic pattern of song variation was consistent with convergent character displacement, with the acoustic structure of *H. peruviana* and *H. subflava* songs being more similar in sympatric taxa than in allopatric taxa (figure 1, S3–S5). Specifically, we found that sympatric lineages *H. p. peruviana* and *H. s. collinsi* differed significantly in only two of seven PCs extracted from song features, as well as in pace (*P* = 0.012), while allopatric lineages *H. p. saturata* and *H. s. subflava* differed in six out of seven PCs and pace (*P* < 0.0001). Strikingly, within one species, songs of *H. s. collinsi* were much more similar to songs of a heterospecific lineage in sympatry (*H. p. peruviana*) than to a conspecific lineage in allopatry (*H. s. subflava*), from which it also differed in six out of seven PCs, and both peak frequency (*P* = 0.0005) and pace (*P* < 0.0001). Song differences between *H. p. peruviana* and *H. p. saturata* are smaller than within *H. subflava* (also differing in two out of seven PCs, and in peak frequency; *P* = 0.0005), consistent with their shallower genetic divergence (table 1, figure 1, S3–S5, ANOVA results: table S5). Nevertheless, by modelling song structure in *H. peruviana* across its range, we found evidence that songs became gradually more similar to those of *H. subflava* in peak frequency (figure 2), as well as in some temporal features, towards the zone of sympatry.

**Table 1.**
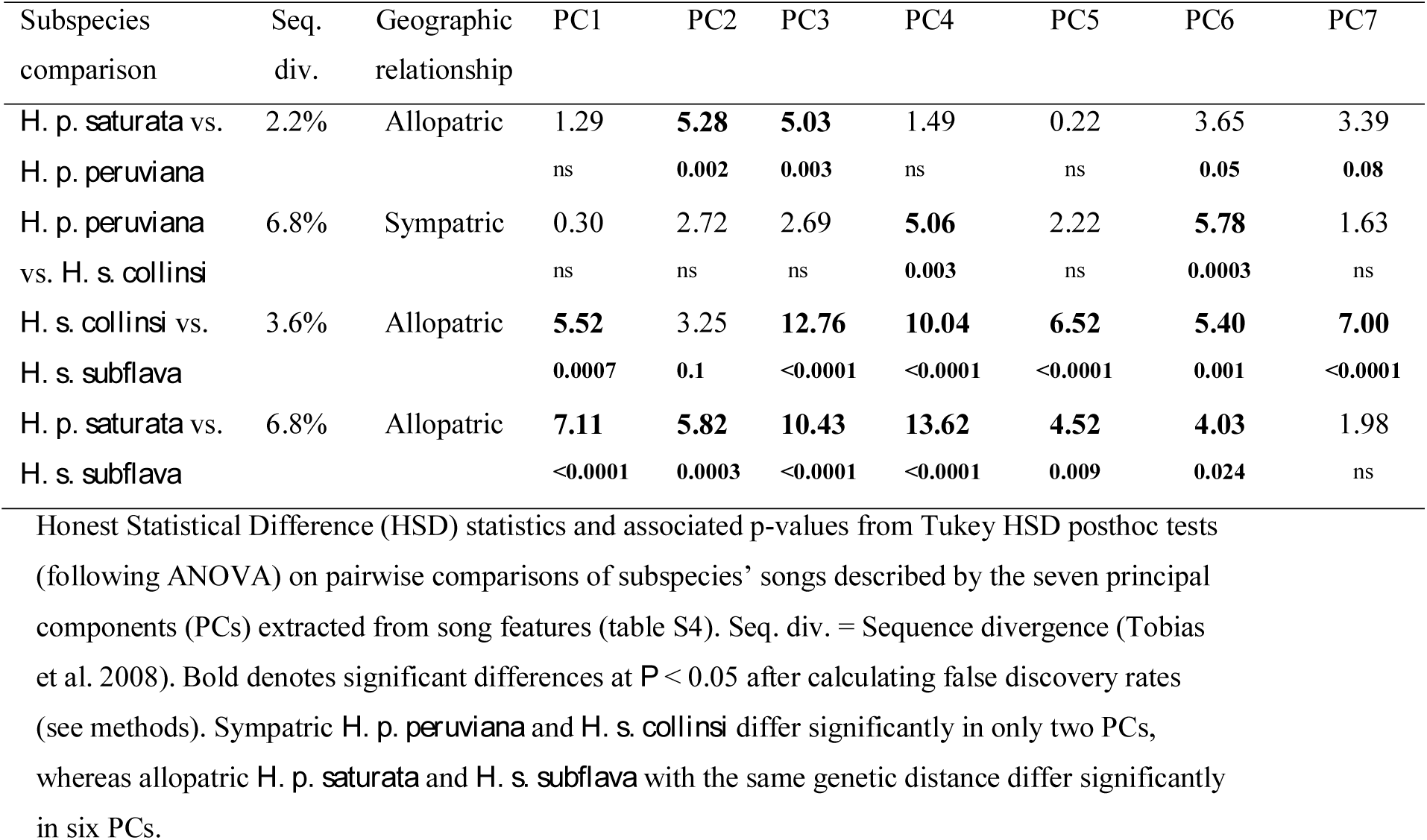
Comparison of song structure in *Hypocnemis* lineages, showing greater signal similarity of sympatric taxa compared to allopatric taxa

| Subspecies comparison | Seq. div. | Geographic relationship | PC1 | PC2 | PC3 | PC4 | PC5 | PC6 | PC7 |
| --- | --- | --- | --- | --- | --- | --- | --- | --- | --- |
| *H. p. saturata* vs. *H. p. peruviana* | 2.2% | Allopatric | 1.29 | **5.28** | **5.03** | 1.49 | 0.22 | 3.65 | 3.39 |
|  |  |  | ns | **0.002** | **0.003** | ns | ns | **0.05** | **0.08** |
| *H. p. peruviana* vs. *H. s. collinsi* | 6.8% | Sympatric | 0.30 | 2.72 | 2.69 | **5.06** | 2.22 | **5.78** | 1.63 |
|  |  |  | ns | ns | ns | **0.003** | ns | **0.0003** | ns |
| *H. s. collinsi* vs. *H. s. subflava* | 3.6% | Allopatric | **5.52** | 3.25 | **12.76** | **10.04** | **6.52** | **5.40** | **7.00** |
|  |  |  | **0.0007** | **0.1** | **<0.0001** | **<0.0001** | **<0.0001** | **0.001** | **<0.0001** |
| *H. p. saturata* vs. *H. s. subflava* | 6.8% | Allopatric | **7.11** | **5.82** | **10.43** | **13.62** | **4.52** | **4.03** | 1.98 |
|  |  |  | **<0.0001** | **0.0003** | **<0.0001** | **<0.0001** | **0.009** | **0.024** | ns |

Honest Statistical Difference (HSD) statistics and associated p-values from Tukey HSD posthoc tests (following ANOVA) on pairwise comparisons of subspecies' songs described by the seven principal components (PCs) extracted from song features (table S4). Seq. div. = Sequence divergence (Tobias et al. 2008). Bold denotes significant differences at $P < 0.05$ after calculating false discovery rates (see methods). Sympatric *H. p. peruviana* and *H. s. collinsi* differ significantly in only two PCs, whereas allopatric *H. p. saturata* and *H. s. subflava* with the same genetic distance differ significantly in six PCs.

**Figure 2.**
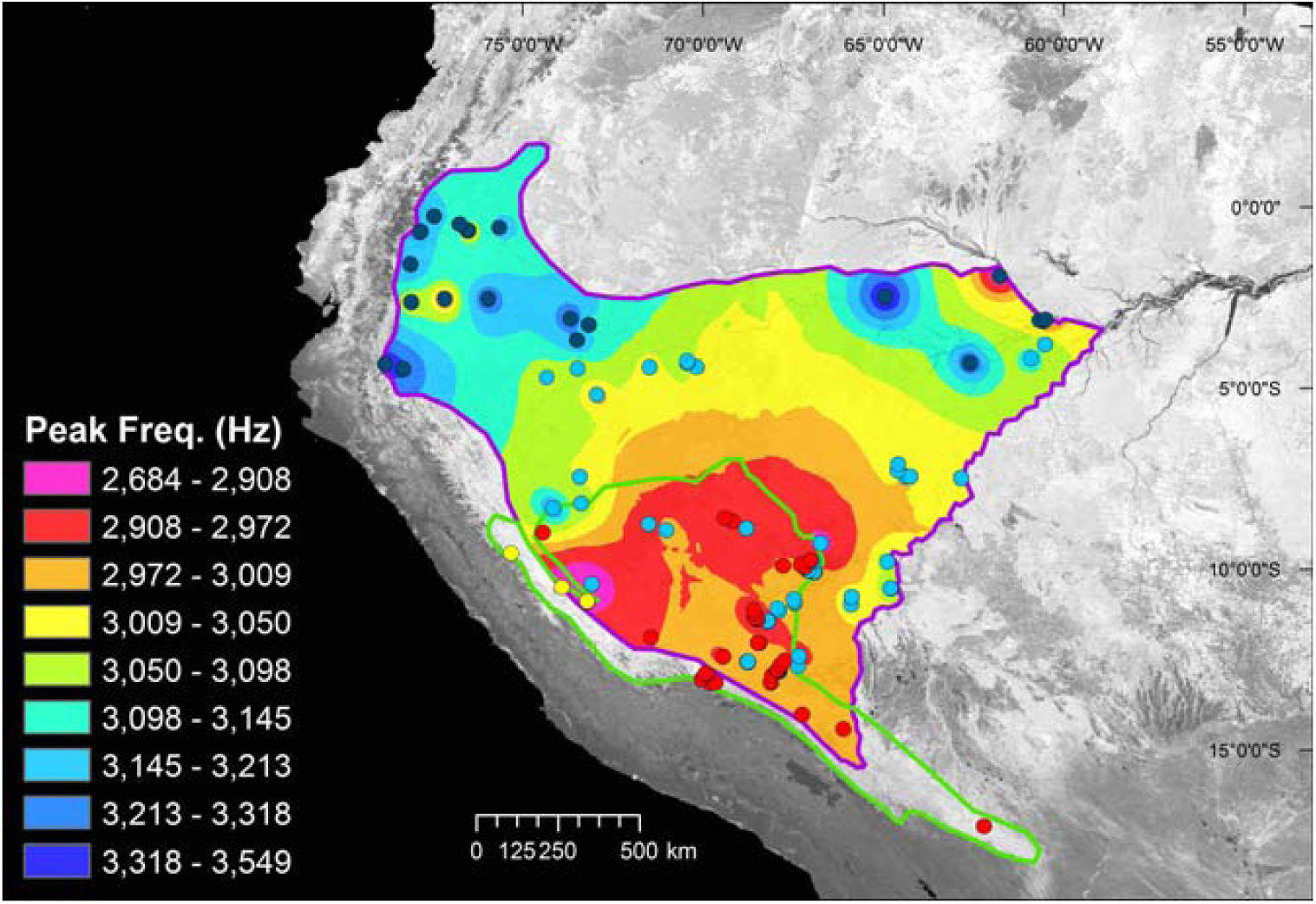
Spatial distribution of peak frequency of the songs of *H. peruviana*, revealing convergence towards the contact zone with *H. subflava*. Shown are the ranges of *H. peruviana* (purple outline), and *H. subflava* (green outline), and their extensive region of overlap. In the song frequency legend, red represents the frequency bandwidth that includes *H. s. collinsi* mean peak frequency of 2916 Hz (mean *H. s. subflava* frequency was 3112 Hz). Frequencies become more different from *H. s. collinsi* as colours diverge along a gradient from pink (lowest peak frequency) to dark blue (highest peak frequency) with greater distance from the contact zone. The most similar peak frequencies (red) are within the contact zone and immediately adjacent areas. The background image illustrates EVI for August 2011. Dots indicate recording localities for *H. p. peruviana* (cyan), *H. p. saturata* (blue), *H. s. subflava* (yellow), and *H. s. collinsi* (red).

We found a significant effect of distance from the contact zone on four PCs. Two of these showed a pattern of convergence: PC6 (GLMM: *z* = 3.26, *P* = 0.001) and PC7 (*z* = 4.27, *P* < 0.0001; table S3). PC6 is associated with mean minimum and peak frequencies; PC7 is associated with the mean and variance in note peak time, a measure of the temporal patterning of notes (Tobias et al. 2014). In other words, the songs of *H. peruviana* became more similar to those of *H. subflava* in pitch and aspects of temporal structure as they approached the contact zone. By contrast, an opposing pattern of divergence towards the contact zone was found in both PC2 (*z* = 2.95, *P* = 0.003) and PC3 (*z* = 2.96, *P* = 0.003). PC2 is associated with mean and variance in bandwidth and mean maximum frequency; PC3 is associated positively with song pace, and pace of first tercile, and negatively with mean and variance in internote intervals. The remaining PCs showed no pattern of convergence or divergence in *H. peruviana* song towards the contact zone (table S6; all results after correcting for false discovery rates (Benjamini and Hochberg 1995, Benjamini and Hochberg 2000, Verhoeven et al. 2005)). Furthermore, we found a significant pattern of convergence towards the contact zone in peak frequency (figure 2, 3*a*), but no significant effect on pace (figure 3*b*, S6, table S6).

**Figure 3.**
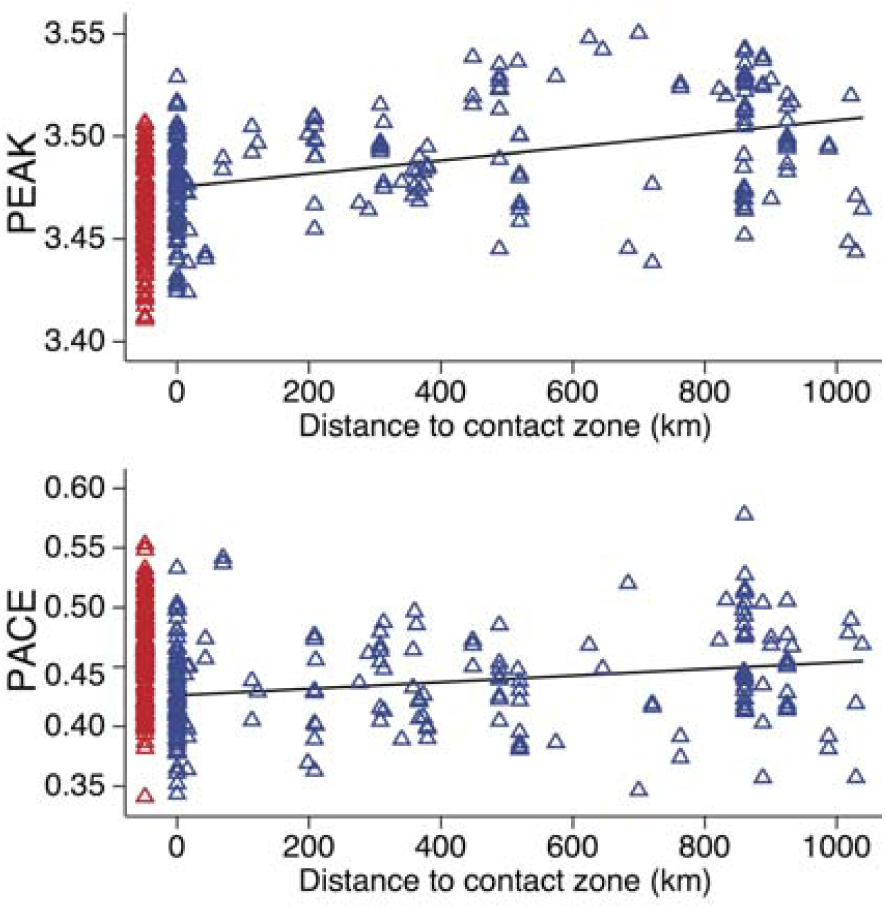
Convergence in song pitch towards the contact zone between *H. peruviana* and *H. s. collinsi*. Shown is log mean peak frequency (a) and log mean pace (b) for each individual of *H. peruviana* (blue) and *H. s. collinsi* (red), plotted against distance from the contact zone, with sympatric individuals illustrated at zero distance.

Incorporating five environmental variables obtained from remote sensing data (table S2) into GLMMs, and after correcting for false discovery rates (table S7), we only found weak relationships between environment and song. Specifically, we found a negative correlation between annual mean temperature and PC7 (*z* = −2.91, *P* = 0.004), and a positive correlation between elevation and the pace-related PC3 (*z* = 3.54, *P* < 0.001. Our overall models include these effects, indicating that overall patterns of song convergence towards the contact zone are not explained by correlations with environmental variables linked to vegetation density. This result is consistent with direct observations at field sites suggesting that vegetation structure of rainforest habitat occupied by *H. peruviana* is roughly similar across its geographical range in Western Amazonia.

## Discussion

Our spatial analyses show that key aspects of the territorial songs of two antbird species converge over large geographical scales from allopatry to sympatry, and also suggest that this pattern arises independently of underlying environmental gradients. In addition, the details of the *Hypocnemis* system allow us to reject two further hypotheses for spatial convergence: heterospecific song copying and hybridisation in sympatry (Tobias and Seddon 2009, Tobias et al. 2012). Evidence for spatial convergence is consistent with evidence of variation among taxa given that *H. s. collinsi* songs are more similar to those of *H. p. peruviana* in sympatry than they are to the songs of allopatric conspecifics (*H. s. subflava*). This finding contrasts with the dominant view that song variation largely reflects phylogeny (Price and Lanyon 2002, Farnsworth and Lovette 2008, Nwankwo et al. 2018), and suggests that interspecific interactions have driven convergent signal evolution in some *Hypocnemis* antbird lineages.

We found some evidence that different aspects of song design diverged towards the contact zone, potentially reflecting divergent selection on particular song traits. However, this spatial pattern is subtle and has not resulted in any significant differences in these songs traits between sympatric subspecies (*H. p. peruviana* and *H. s. collinsi*). Thus, the strongest and most consistent spatial and taxonomic pattern we detected was song convergence in both our study species. This pattern is opposite to the predictions of standard character displacement theory (Servedio 2004, Pfennig and Pfennig 2009), but consistent with theoretical models of convergent ACD mediated by social competition (Cody 1969, Grether et al. 2009, Grether et al. 2017).

One aspect of our results that deviates from the predictions of convergent ACD is that signal convergence appears to be asymmetric, with song characters in sympatry sometimes non-intermediate between the allopatric means (figure 1) or occasionally overshooting the respective mean trait in heterospecifics (figure 3). Only one trait (PC1) shows the symmetrical pattern of meeting in the middle when lineages interact. It is not clear why asymmetric convergence arises. It may be because of asymmetries in competitive ability, or simply because characters involved in social competition evolve rapidly (West-Eberhard 1983) and at different rates across lineages. Signal design may also be shaped by interactions with further species, perhaps through signal partitioning to minimise interference (Chek et al. 2003). However, previous community-wide analyses in the sympatric zone found no evidence of partitioning across 307 bird species (Tobias et al. 2014). Moreover, playback experiments in their contact zone indicate that neither *H. p. peruviana* nor *H. s. collinsi* respond to the (highly divergent) song of their closest sympatric relative (*Drymophila devillei*) (Tobias and Seddon 2009). Although minor asymmetries in trait variation remain unexplained in this system, key traits are nonetheless more similar in sympatry than allopatry, and thus both spatial and taxonomic analyses provide more support for convergent ACD than divergent RCD.

Classic examples of song divergence in sympatry are widely reported (Kirschel et al. 2009, Lemmon 2009, Grant and Grant 2010) and intuitively plausible given that divergent signals are thought to reduce the costs of competitive or reproductive interference (Groning and Hochkirch 2008). Why might divergent selection be outweighed by convergence in *Hypocnemis*? Like most birdsongs (Collins 2004), *Hypocnemis* songs function in both mate attraction and territory defence (Tobias et al. 2011), yet interference costs appear to be circumvented in this case by secondary adaptations, including divergent visual signals and finely tuned receiver perception (Tobias and Seddon 2009, Seddon and Tobias 2010). In particular, previous experiments have shown that females of both species can easily differentiate between conspecific and heterospecific male songs, suggesting that accurate mate recognition can accommodate convergent ACD in song characters (Seddon and Tobias 2010). It seems likely that selection for mate recognition targets particular components of song structure in *Hypocnemis*, perhaps explaining why some acoustic characters of *H. peruviana* and *H. subflava* appear divergent, or at least non-convergent, in sympatry.

In *H. peruviana*, our data reveal that convergence is gradual from allopatry towards the contact zone, rather than a rapid step in characters at the boundary between allopatry and sympatry. We suspect that patterns of gradual signal convergence extending beyond the contact zone may be commonplace for two reasons. First, geographical ranges fluctuate, and thus the boundary between sympatry and allopatry may shift over time, blurring any abrupt spatial step in signal traits or their underlying genes. Second, signal traits under selection in the contact zone may be transferred into allopatric populations in the absence of barriers to gene flow, producing a gradient of song characters.

A possible alternative explanation for spatial gradients in song is that they are driven by an underlying environmental gradient coupled with acoustic adaptation (e.g., Slabbekoorn and Smith 2002, Seddon 2005, Kirschel et al. 2009, Kirschel et al. 2011). Because we sampled songs from numerous localities, and often from digital sound archives, we were not able to directly estimate variation in sound transmission or vegetation density, both of which have been shown to shape acoustic signal structure (Wiley and Richards 1982, Slabbekoorn and Smith 2002, Kirschel et al. 2009, Tobias et al. 2010, Kirschel et al. 2011). Instead, we estimated environmental conditions using remote sensing data, representing variation in vegetation density, elevation and climate. Although there are limitations to remote sensing data because of the relative coarseness of sampling, the variables we selected are widely used to control for environmental variation in studies of signal evolution (Kirschel et al. 2009, Kirschel et al. 2009, Kirschel et al. 2011, Smith et al. 2013). Moreover, the levels of environmental variation we detected accurately reflect the fact that the global range of *H. peruviana* is almost entirely restricted to lowland tropical rainforest with little variation in topography or vegetation density (Isler et al. 2007, Tobias and Seddon 2009). Our analyses therefore suggest that convergence in song traits does not arise through ecological selection exerted by the signal transmission properties of the environment, but rather by a socially mediated mechanism such as ACD.

Convergent ACD may be more frequent in diverse communities, including tropical rainforests, where interspecific territoriality is widespread (Robinson and Terborgh 1995), and many species coexist with similar ecological and signalling traits (Robinson and Terborgh 1995, Tobias et al. 2014). Regardless of whether species in hyper-diverse communities have assembled through ecological sorting of existing phenotypes (Pigot et al. 2016), or instead converged in ecological characters towards peaks in the adaptive landscape (Scheffer and van Nes 2006), their coexistence may be facilitated by competitor recognition (Grether et al. 2009, Grether et al. 2017). From this perspective, convergent ACD may provide further insight into the mechanisms explaining patterns of species coexistence and the build-up of tropical diversity in vertebrate systems.

Divergent forms of character displacement, including both RCD and ACD, have traditionally been thought to play an important role in driving phenotypic divergence and explaining signal variation across geographic space (Brown and Wilson 1956, Schluter et al. 1985, Kirschel et al. 2009). While these patterns may indeed be widespread, the allopatry-to-sympatry gradients explored in this study provide new evidence that ACD can also shape signal phenotypes the opposite way by driving broad-scale convergence towards regions of coexistence between ecological competitors. Our results add weight to previous theoretical and local-scale evidence for convergent ACD (Cody 1969, Grether et al. 2009, Tobias and Seddon 2009, Grether et al. 2017), and highlight its role in shaping geographical variation of animal signals at continental scales.

## Acknowledgements

We are grateful to numerous field ornithologists for contributing songs to this study, particularly M and P Isler and G Budney (Macaulay Library, Cornell). We also thank A Aleixo, R Amable, M Bianchini, E Guilherme, A Jameson, A Lees, P Long, H MacGregor, B Nelson, E Nishikawa, G Olah, B Poje, D Romo, T Valqui, N Yavit, plus the staff at Amazon Conservation Association, Los Amigos Research Station (CICRA), Amazon Research Conservation Center, Heath River Wildlife Center, Tiputini Biodiversity Station, and Rainforest Expeditions for assistance with data collection, logistics and analyses. This research was supported by an FP7 Marie Curie International Incoming Fellowship (ANGK), a Royal Society University Research Fellowship (NS), and a John Fell award (JAT).

