## Supplementary material for "Range-wide spatial mapping reveals convergent character displacement of bird song"

ELECTRONIC SUPPLEMENTARY MATERIAL

**Contents**

Expanded methods: Song sampling, Song processing, Quantifying environmental gradients, Inverse distance weighting

Supplementary Figures (S1–S7)

Data Tables (S1–S3)

Statistical Tables (S4 - S6)

**Expanded methods**

**Song sampling**

We compiled sound files from original field recordings and several different digital sound archives. This approach means that the sound files used in our analyses were recorded by different observers and with different recording devices, raising the possibility that our results may be influenced by underlying biases, for instance if particular file types or recording devices are biased towards sympatric or allopatric samples. However, recent analyses have shown that compressed MP3 files do not lead to any systematic deviations in commonly used acoustic measurements [1]. Minor problems can arise with extreme value parameters, so we adopted the recommended processing pipeline to minimize these distortions [1]. Moreover, the final sample includes 353 uncompressed recordings and only 2 compressed MP3s, suggesting that file type is unlikely to influence our results.

The role of different recording devices in biasing acoustic structure is harder to pinpoint, particularly as recording device is often not specified in archived material. However, available data suggests that there was no spatial bias in the use of recording devices between sympatry and allopatry. Of 355 recordings used in total, 311 (87.6%) were by recordists recording in both sympatry and allopatry. Only 44 cuts were obtained by recordists active only in sympatry or in allopatry, with no more than five recordings attributed to any one of those recordists. Of course, many of these allopatry-only or sympatry-only recordists will use devices that are the same as each other, or the same as the six major contributors: A. Kirschel (123 recordings), N. Seddon and J. Tobias (71 recordings), B. Whitney (50 recordings), K. Zimmer (26 recordings), the late T. Parker (19 recordings) and a group of recordists from Cornell on expedition using the same recording equipment, B. Winger, M. Harvey, G. Seeholzer, and R. Terrill (12 recordings). These six groups all recorded songs in both sympatry and allopatry, suggesting that biases in recording equipment do not influence our results. In any case, previous studies have not highlighted a problem with pooling data across multiple high-quality recording devices, and indeed this is now the standard approach in many recent bioacoustic studies at regional or global scales [2-6].

**Song processing**

Song processing methods followed previous studies in suboscine passerines [5, 7]. All songs were filtered using a 10^th^ order highpass Butterworth filter (cut-off frequency = 400 Hz) before final broadband spectrograms were generated (window = Hann, bandwidth = 256 Hz, Fast Fourier Transform = 1024, overlap = 0.875). Spectrograms were visualized with a custom graphical user interface (GUI) and manually segmented using on-screen cursors to record sample number at note onset and offset. A note was defined as a continuous trace on the spectrogram with amplitude much greater than that of background noise. We used 10% of peak energy as a threshold for the temporal extent of a note. Segmented songs were then analyzed using a custom MatLab script code [5]. All MP3 files were converted to wav for analyses. To minimize the effects of extreme value parameters resulting from compression, we restricted our analyses to measures averaged over the entire song, or for five pace measures averaged over terciles of the entire song for comparison purposes, following published recommendations [1].

**Quantifying environmental gradients**

Where possible, we mapped exact localities in ArcGIS 10.1 of individual territories using GPS coordinates of recordings collected during fieldwork. Other coordinates were provided with recordings obtained from other sources (xeno-canto (<http://xeno-canto.org>), Macaulay Library (<http://macaulaylibrary.org>), private collections), and for a few remaining sites, were estimated from Google Earth. In these cases of indirect spatial sampling, we typically found a single set of coordinates representing several recordings from the same site, meaning that the exact location of each recording was uncertain. To address this issue, we extracted interpolated remote sensing data for a 5 x 5 pixel area around the available coordinates, and assigned these data to the songs recorded at that locality.

In order to delimit the contact zone, we plotted all the coordinates from our recordings and those obtained from other sources. Sites where both species were recorded were defined as sympatric. We examined how the distribution of these sites compared with illustrations of the contact zone in previous publications [7, 8]. Parts of Western Acre, Brazil, and Ucayali, Peru, remain very inaccessible and poorly known, and thus the known extent of the contact zone is probably an underestimate. Because it seems likely that both species are present throughout the extent of *Guadua* bamboo (the preferred habitat of *H. s. collinsi*), we used the limit of *Guadua* bamboo distribution [9] to determine the extent of *H. s. collinsi*’s range in these unsurveyed areas.

Most of our song samples were based on recordings made during fieldwork in 2011, during which we visited recording localities across the range of *Hypocnemis peruviana* from allopatry in Ecuador (Jan-Feb 2011) and Brazil (Jul-Aug 2011), to numerous sites within the contact zone. Thus, we based our environmental gradients analysis on data extracted from around that period. For the Enhanced Vegetation Index, we extracted data from a 16-day granule covering a period from August-September 2011 because the finer temporal resolution specificity allowed us to relate vegetation densities to the period of extensive fieldwork carried out in Brazil, Peru and Bolivia between Jul and Nov 2011. Percent tree cover data were obtained for the previous year (2010) from the vegetation continuous field product (VCF) based on the Moderate Resolution Imaging Spectroradiometer (MODIS; http://modis.gsfc.nasa.gov). We could have used either data for the whole of 2010 or 2011, but chose 2010 as the focal year for tree cover data in order to reduce the possibility of measuring habitat variables from areas deforested after recordings were obtained (some sites were being deforested during surveys in 2011 - thus tree cover data for the whole of 2011 may have been inaccurate).

A number of song recordings were collected during earlier surveys between 1975 and 2000. Extracting environmental information from those specific years was not possible because MODIS data are only available from 2000 onwards. Because we could not obtain environment data for all recordings, we did not attempt to obtain further environment data specific to recording dates between 2000 and 2010. Instead, we used vegetation structure data extracted from the same periods as above (2010 for VCF; Aug–Sep 2011 for EVI) to maximise consistency. This is unlikely to generate inaccurate measures as habitat has remained fairly constant at most study sites during the study period, and the error introduced by any local-scale vegetation change is reduced by extracting interpolated environmental data from a 5 x 5 pixel area for site coordinates.

The WorldClim data [10] from which we extracted two bioclimatic variables (annual mean temperature and annual precipitation) are based on records from the period 1950–2000. While these data do not cover the entire study period, they are nonetheless representative of mean climatic differences across the study region. Elevation data were extracted from 2010.

**Inverse Distance Weighting (IDW)**

IDW predicts the value for an unmeasured location using a set of measured values around it, with values closer to the predicted location having more influence than more distant. values (ArcGIS Help 10.1). We set the parameters in ArcGIS to use the 50 geographically closest measurements of songs in the interpolation of each cell value. Using a higher number of nearby measurements controls for possible outliers in nearby sites and for those areas that are poorly sampled. Selecting fewer points for the interpolation resulted in outliers having larger apparent effects on the distribution of song variation across the landscape. We opted for a power value of 1.5, to allow for surrounding points at greater distances to have a little more influence on each cell value than with the default setting of 2 (and within the ArcGIS recommended range of 0.5–3), because the extent of geographic sampling varied across the range of the species, with tighter sampling in sympatry than in some areas in allopatry. We chose the natural breaks [11] option to delineate nine peak frequency ranges (see Fig. 2, S6).

**Supporting Figures**

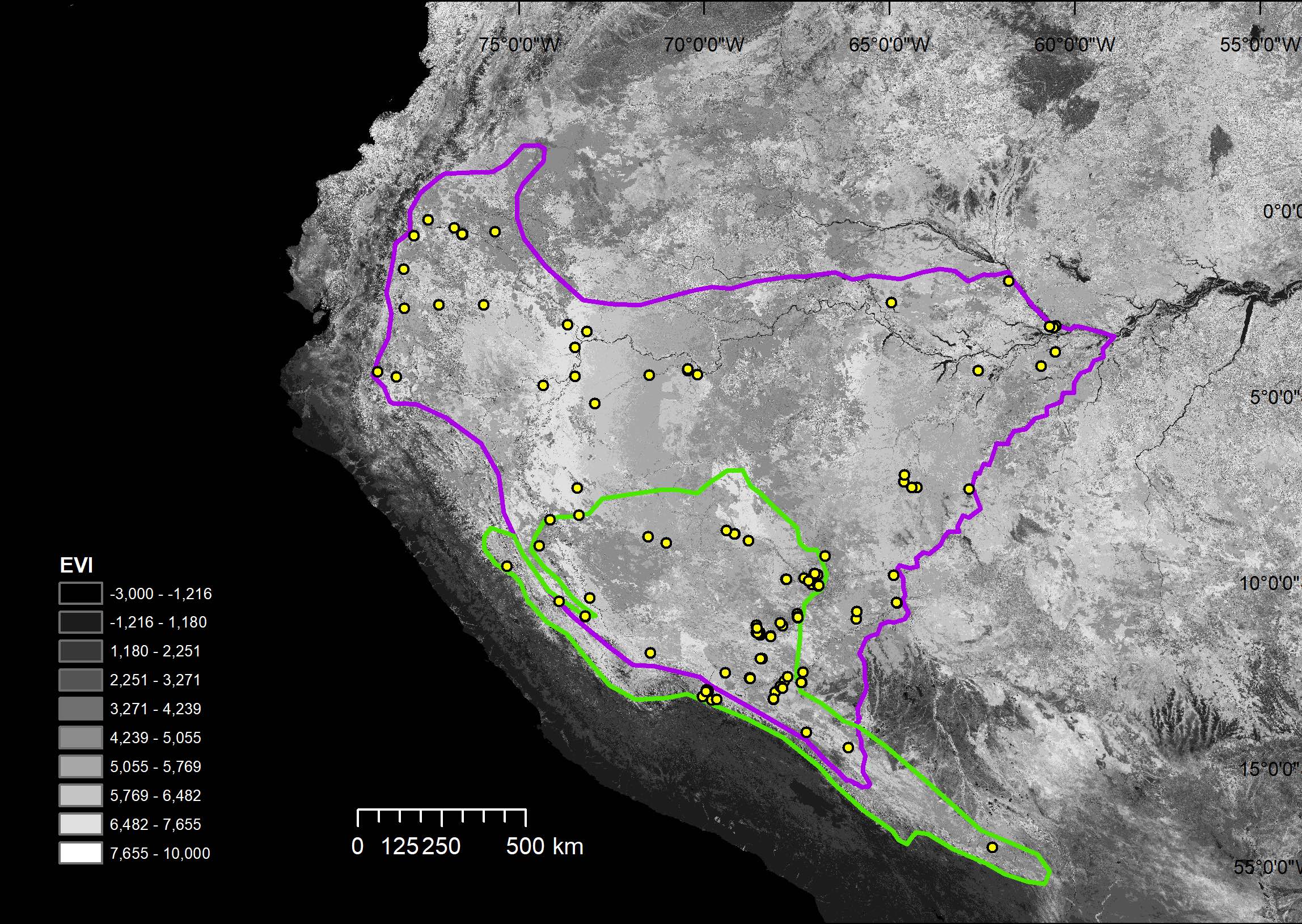

**Figure S1** Enhanced vegetation index (EVI) raster for August 2011 overlaid by the ranges of *H. peruviana* (purple outline), and *H. subflava* (green outline), with the contact zone occurring where they overlap. The legend divides EVI into 10 categories separated by natural jenks, with brighter shades representing more densely vegetated areas. Recording localities are shown as yellow dots.

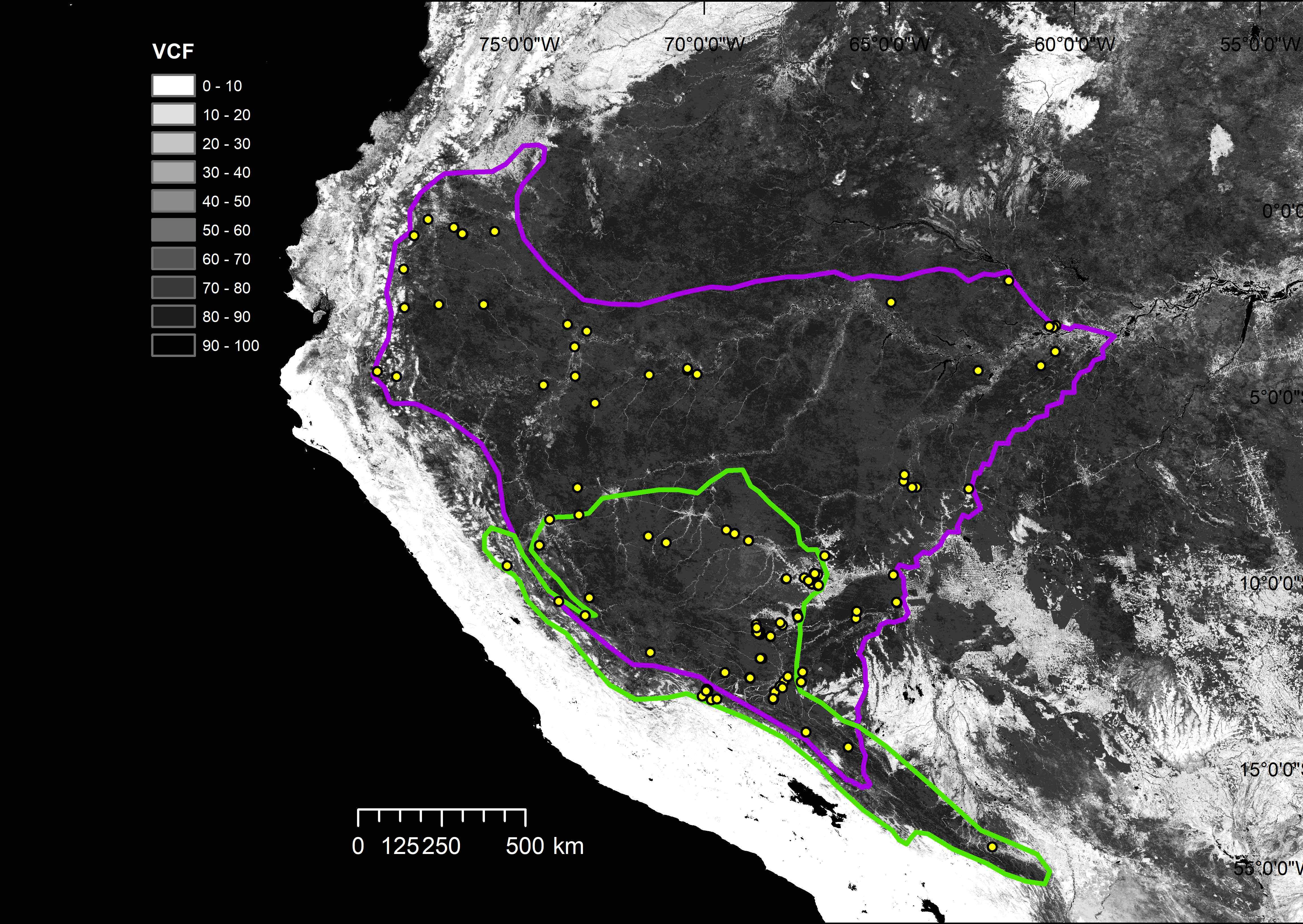

**Figure S2** Vegetation continuous field (VCF) raster for 2010, representing percent tree cover, overlaid by the ranges of *H. peruviana* (purple outline), and *H. subflava* (green outline), with the contact zone occurring where they overlap. The legend divides VCF into 10 equal percent tree cover categories, with darker shades representing higher densities. Recording localities are shown as yellow dots.

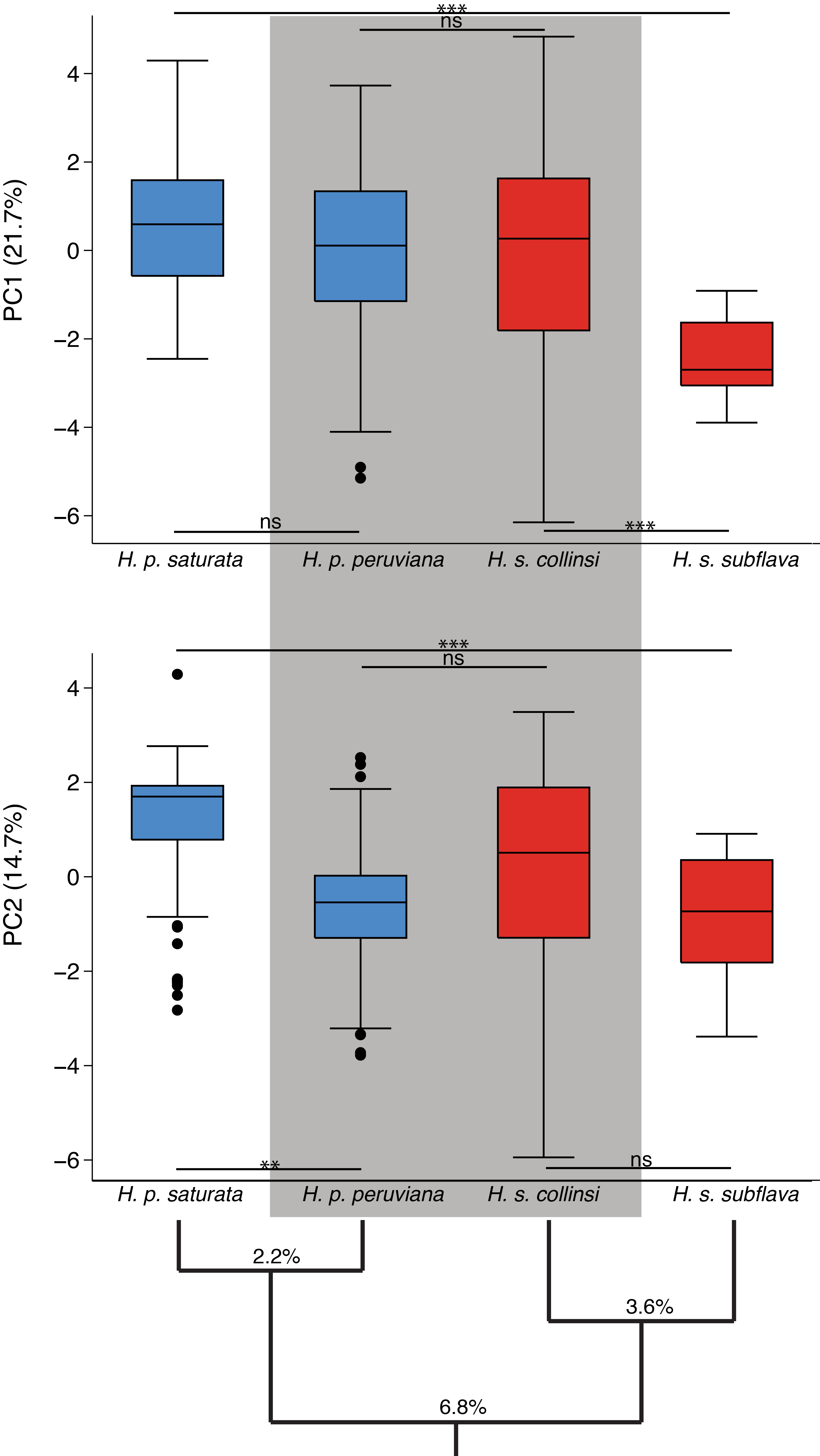

**Figure S3** Songs of *H. p. peruviana* and *H. s. collinsi* (shaded in gray; partly sympatric) are more similar in PC1 than *H. s. collinsi* is to conspecific *H. s. subflava*, and thus more similar to each other than would be expected based on their genetic distance (shown as % mtDNA sequence divergence). PC2 also shows greater similarity between partly sympatric forms than between *H. p. peruviana* and conspecific *H. p. saturata* (allopatric). The two partly sympatric forms are thus more similar than would be expect based on their genetic distance, while each differs significantly from much closer related conspecifics in one of the two PCs. Levels of significance are following FDR correction: * P < 0.05, ** P < 0.01, *** P < 0.001, ns = not significant.

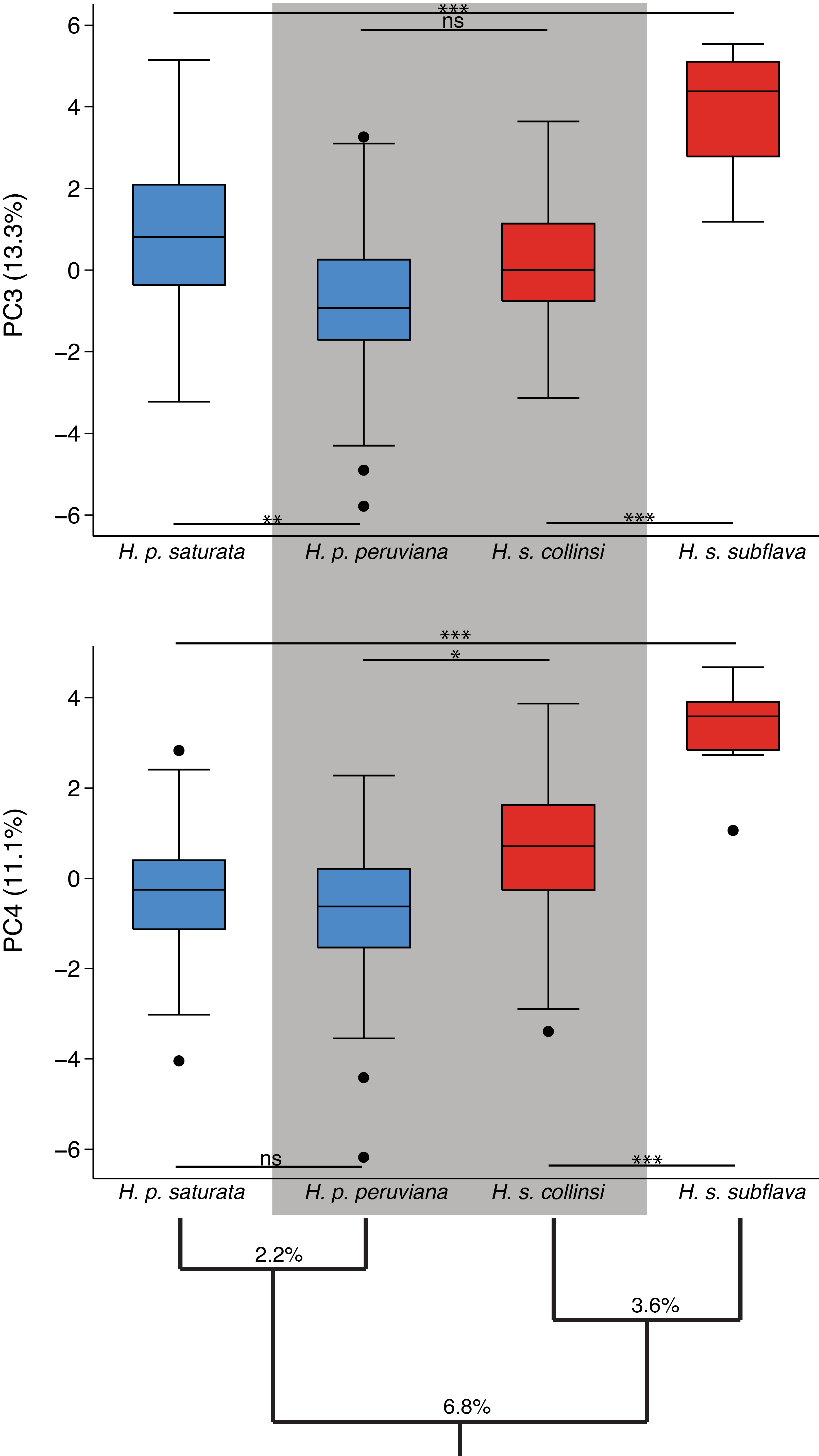

**Figure S4** Songs of *H. p. peruviana* and *H. s. collinsi* (partly sympatric) are more similar in both PC3 and PC4 than *H. s. collinsi* is to conspecific *H. s. subflava*, consistent with the hypothesis that *H. s. collinsi* has converged towards *H. p. peruviana* and away from *H. s. subflava* in song traits represented by these PCs. *H. p. peruviana* is most similar to allopatric conspecific *H. p. saturata* in PC4, but most similar to partly sympatric heterospecific *H. s. collinsi* in PC3 (cladogram shows % mtDNA sequence divergence). Levels of significance are following FDR correction: * P < 0.05, ** P < 0.01, *** P < 0.001, ns = not significant.

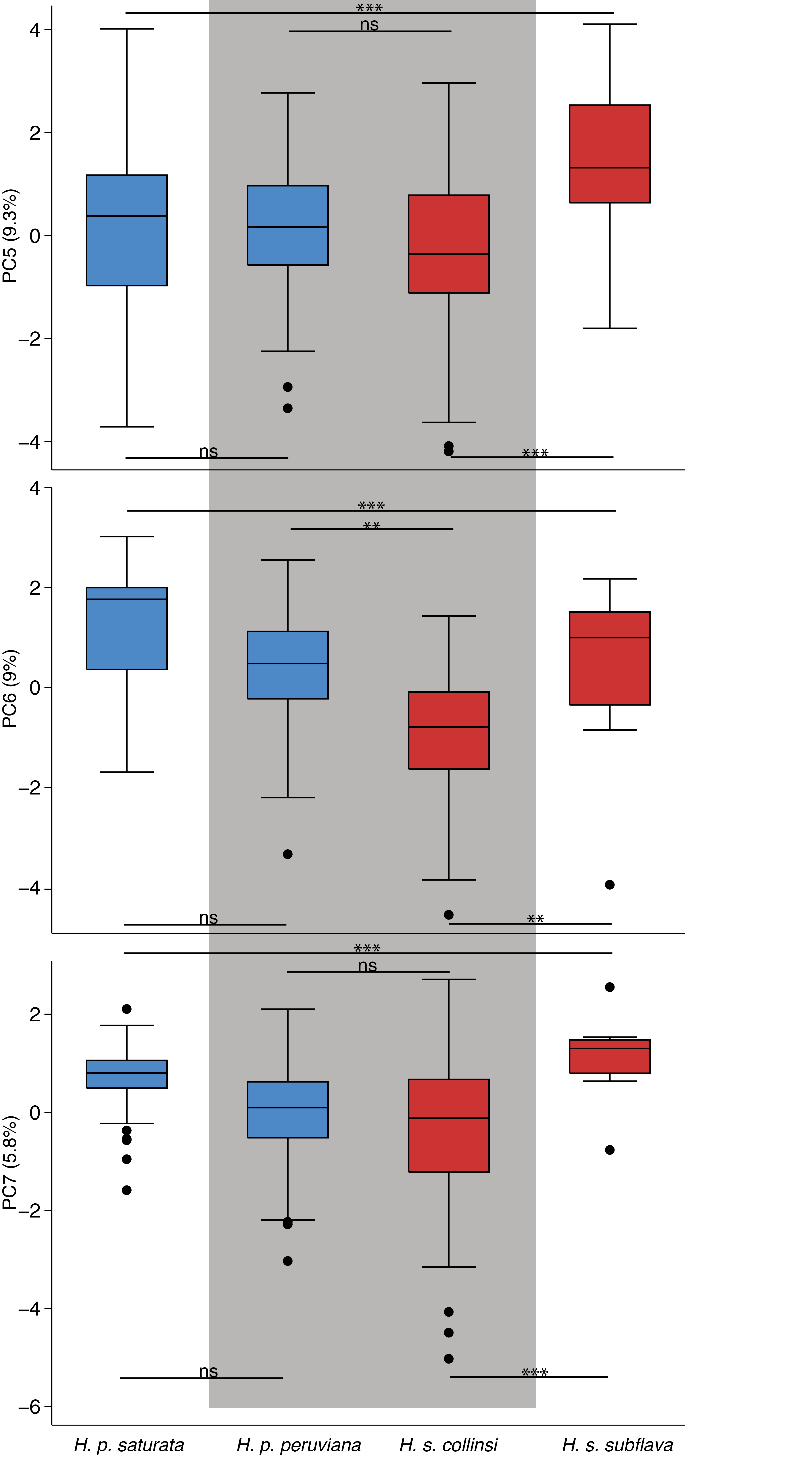

**Figure S5** Songs of *H. p. peruviana* and *H. s. collinsi* (partly sympatric) differ significantly in PC6, and *H. p. peruviana* song appears to converge towards *H. s. collinsi* and away from *H. p. saturata* (allopatric). Differences between heterospecific *H. p. peruviana* and *H. s. collinsi* (partly sympatric) in PC5 and PC7 are smaller than those between conspecific *H. s. collinsi* and *H. s. subflava* (allopatric), thus contrasting patterns of genetic distance (shown as % mtDNA sequence divergence). Levels of significance are following FDR correction: * P < 0.05, ** P < 0.01, *** P < 0.001, ns = not significant.

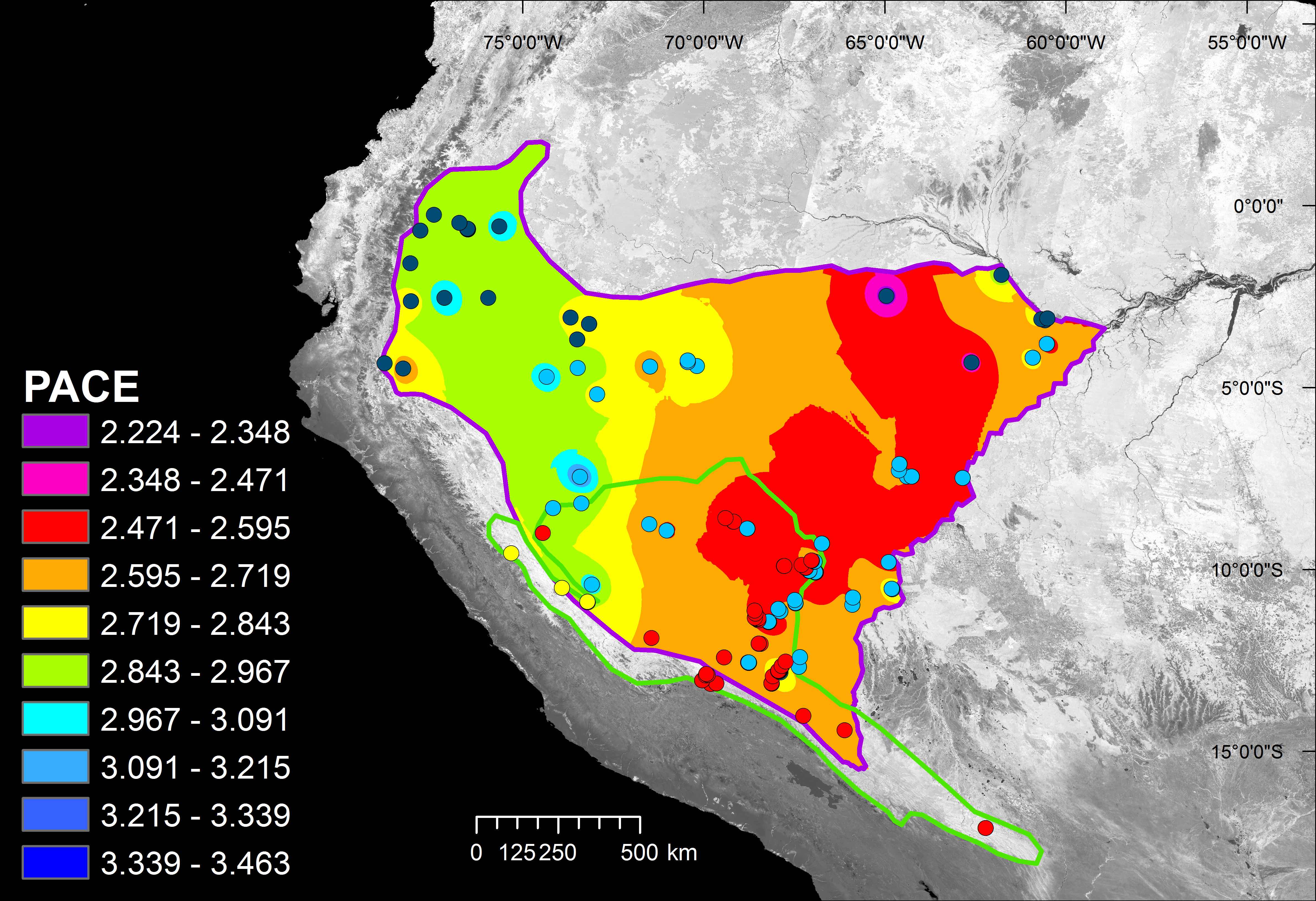

**Figure S6** Spatial distribution of the overall pace of the songs of *H. peruviana*, revealing no divergence or convergence towards the contact zone with *H. subflava*. Shown are the ranges of *H. peruviana* (purple outline), and *H. subflava* (green outline), with the contact zone occurring where they overlap. In the song pace legend, green represents the pace range that includes *H. s. collinsi* mean pace of 2.91 notes per second. The background image illustrates the Enhanced Vegetation Index for August 2011 (representing canopy structural variation). Recording localities are shown as dots for *H. p. peruviana* (cyan), *H. p. saturata* (blue), *H. s. subflava* (yellow), and sites for *H. s. collinsi* where *H. peruviana* was not recorded (red).

**Table S1** Location and sampling of study sites

| Site | Country | Latitude | Longitude | Subspecies |
| --- | --- | --- | --- | --- |
| Coca, 34 km N of; Napo | Ecuador | -0.25 | -77.07 | *saturata* (1) |
| La Selva Lodge,Napo | Ecuador | -0.47 | -76.37 | *saturata* (4) |
| Imuya Cocha; Napo | Ecuador | -0.57 | -75.28 | *saturata* (1) |
| Tiputini Biodiversity Station | Ecuador | -0.64 | -76.15 | *saturata* (27) |
| Loreto, 15 to 80 km W by road; Napo | Ecuador | -0.68 | -77.45 | *saturata* (2) |
| Canelos; Pastaza | Ecuador | -1.58 | -77.75 | *saturata* (1) |
| Jau, Parque Nacional de; AM | Brazil | -1.9 | -61.51 | *saturata* (1) |
| Amani, Lago, AM | Brazil | -2.48 | -64.69 | *saturata* (1) |
| El Tigre; Loreto | Peru | -2.53 | -75.65 | *saturata* (1) |
| Kapawi Lodge; Pastaza | Ecuador | -2.53 | -76.86 | *saturata* (1) |
| Miazal, Morona-Santiago | Ecuador | -2.62 | -77.78 | *saturata* (2) |
| Libertad, 1.5 km S; Loreto | Peru | -3.07 | -73.42 | *saturata* (1) |
| Jacare Un Bau, AM | Brazil | -3.1 | -60.3 | *saturata* (1) |
| Sao Francisco, Fazenda, AM | Brazil | -3.12 | -60.47 | *saturata* (1) |
| Pousada Amazonia, AM | Brazil | -3.15 | -60.37 | *saturata* (2) |
| Sucusari, Quebrada; Loreto | Peru | -3.25 | -72.92 | *saturata* (8) |
| San Andres, Loreto | Peru | -3.67 | -73.28 | *saturata* (3) |
| Careiro do Castanho; AM | Brazil | -3.8 | -60.36 | *peruviana* (2) |
| Tupana | Brazil | -4.18 | -60.77 | *peruviana* (10) |
| Santa Rita, Loreto | Peru | -4.25 | -70.29 | *peruviana* (3) |
| Palmari, Reserva Natural; AM | Brazil | -4.3 | -70.29 | *peruviana* (6) |
| Macaca, Igarape do, AM | Brazil | -4.31 | -62.47 | *saturata* (2) |
| Miazi; Zamora-Chinchipe | Ecuador | -4.33 | -78.65 | *saturata* (1) |
| Benjamin Constant, AM | Brazil | -4.41 | -70.05 | *peruviana* (1) |
| Limera, Quebrada; Loreto | Peru | -4.42 | -71.34 | *peruviana* (1) |
| Yarapa Reserve; Loreto | Peru | -4.46 | -73.34 | *peruviana* (1) |
| Huampami; AM | Peru | -4.47 | -78.15 | *saturata* (1) |
| Samiria, Rio; Loreto | Peru | -4.7 | -74.22 | *peruviana* (1) |
| Angamos, Colonia; Loreto | Peru | -5.18 | -72.88 | *peruviana* (1) |
| Livramento, Igarape; AM | Brazil | -7.1 | -64.77 | *peruviana* (2) |
| Labrea, AM | Brazil | -7.28 | -64.82 | *peruviana* (2) |
| Labrea, 24 km E, AM | Brazil | -7.44 | -64.62 | *peruviana* (3) |
| Labrea, 36-42 km E, AM | Brazil | -7.44 | -64.49 | *peruviana* (3) |
| Serra do Divisor, Parque Nacional, AC | Brazil | -7.45 | -73.68 | *peruviana* (2) |
| Humaita, AM | Brazil | -7.48 | -63.08 | *peruviana* (8) |
| Abujao; Ucayali | Peru | -8.18 | -73.77 | *peruviana* (2) |
| Yarina-Cocha, Lago; Ucayali | Peru | -8.31 | -74.58 | *collinsi* (1) *peruviana* (3) |
| Manoel Urbano to Feijo, AC | Brazil | -8.67 | -69.67 | *collinsi* (5) |
| Ouro Preto, Igarape, AC | Brazil | -8.76 | -71.99 | *peruviana* (1) |
| Manoel Urbano, AC | Brazil | -8.87 | -69.29 | *collinsi* (2) *peruviana* (2) |
| Seringal Ocidente, AC | Brazil | -8.93 | -71.54 | *peruviana* (1) |
| 35km NE of Tingo Maria; Santa Elena | Peru | -9 | -75 | *collinsi* (1) |
| Boca do Acre road, AM | Brazil | -9.28 | -67.29 | *peruviana* (4) |
| Tingo Maria, 25 km SW; Huanuco | Peru | -9.55 | -76 | *subflava* (1) |
| Humaita Reserve, AC | Brazil | -9.76 | -67.65 | *collinsi* (7) *peruviana* (6) |
| Abuja, Rio; Pando | Bolivia | -9.8 | -65.53 | *peruviana* (1) |
| Bujari, AC | Brazil | -9.88 | -67.97 | *collinsi* (1) |
| Ramal Jarinal, AC | Brazil | -9.9 | -68.47 | *collinsi* (5) |
| Parque Zoobotanico, Rio Branco, AC | Brazil | -9.95 | -67.87 | *collinsi* (5) |
| Parque Chico Mendes, Rio Branco, AC | Brazil | -10.04 | -67.79 | *peruviana* (3) |
| Catuaba Reserve, AC | Brazil | -10.07 | -67.62 | *collinsi* (9) *peruviana* (11) |
| Puntijao | Peru | -10.41 | -73.95 | *peruviana* (3) |
| Prov. Oxapampa; Distrito Puerto Bermudez | Peru | -10.5 | -74.81 | *peruviana* (2) *subflava* (1) |
| Cachuela | Bolivia | -10.53 | -65.59 | *peruviana* (9) |
| San Juan de Nuevo Mundo; Pando | Bolivia | -10.77 | -66.73 | *peruviana* (2) |
| 13km SE of Oventeni; Monte Tabor | Peru | -10.9 | -74.18 | *subflava* (6) |
| Seringal Cachoeira, AC | Brazil | -10.9 | -68.37 | *collinsi* (1) *peruviana* (3) |
| Ingavi; Pando | Bolivia | -10.97 | -66.79 | *peruviana* (1) |
| Camino Mucden; Pando | Bolivia | -11.08 | -68.89 | *peruviana* (3) |
| Inapari Road | Peru | -11.13 | -69.56 | *collinsi* (1) |
| Cobija, 12-20 km SW; Pando | Bolivia | -11.15 | -68.85 | *peruviana* (2) |
| Belen, Madre de Dios | Peru | -11.22 | -69.57 | *collinsi* (1) |
| Iberia, | Peru | -11.34 | -69.51 | *collinsi* (2) |
| Pacahuara Rd | Peru | -11.35 | -69.58 | *collinsi* (4) |
| Oceania | Peru | -11.4 | -69.52 | *collinsi* (3) |
| Extrema | Bolivia | -11.44 | -69.24 | *collinsi* (3) *peruviana* (4) |
| Provencia La Concepcion; Cuzco | Peru | -11.88 | -72.65 | *collinsi* (1) |
| ARCC, Las Piedras River | Peru | -12.04 | -69.67 | *collinsi* (6) |
| Chive | Bolivia | -12.41 | -68.61 | *peruviana* (1) |
| Manu Wildlife Centre; Madre de Dios | Peru | -12.42 | -70.75 | *collinsi* (4) *peruviana* (2) |
| Cuzco Amazonica Lodge | Peru | -12.53 | -69.05 | *collinsi* (1) |
| CICRA | Peru | -12.56 | -70.1 | *collinsi* (19) *peruviana* (17) |
| Puerto Maldonado | Peru | -12.65 | -69.19 | *collinsi* (6) |
| Heath River Wildlife Center | Bolivia | -12.68 | -68.71 | *peruviana* (2) |
| Posada Amazonas, Tambopata | Peru | -12.8 | -69.31 | *collinsi* (9) *peruviana* (3) |
| Explorer's Inn; Madre de Dios | Peru | -12.83 | -69.28 | *collinsi* (7) *peruviana* (2) |
| Amazonia Lodge; Madre de Dios | Peru | -12.87 | -71.38 | *collinsi* (3) |
| Atalaya and Pilcopata, between; Cuzco | Peru | -12.9 | -71.35 | *collinsi* (2) |
| Patria, Manu Road | Peru | -12.91 | -71.41 | *collinsi* (1) |
| Malinowski CP, Tambopata | Peru | -12.93 | -69.52 | *collinsi* (1) |
| Sandro (Cosnipata valley); Cuzco | Peru | -13.05 | -71.55 | *collinsi* (5) |
| Sandro (Cosnipata valley); Cuzco | Peru | -13.15 | -71.33 | *collinsi* (1) |
| Ccolpa de Guacamayos | Peru | -13.12 | -69.62 | *collinsi* (4) |
| 16km SW of Pilcopata | Peru | -13.13 | -71.17 | *collinsi* (5) |
| Tambopata Research Center | Peru | -13.13 | -69.61 | *collinsi* (6) |
| Curva Alegre; Puno | Peru | -14.02 | -68.97 | *collinsi* (2) |
| Chalalan | Bolivia | -14.42 | -67.92 | *collinsi* (2) |
| Sajta | Bolivia | -17.11 | -64.78 | *collinsi* (15) |

Latitude and longitude are given in decimal degrees.

Sample size of songs recorded (number of individuals) are shown in parentheses after subspecies name.

Details for each recording, including the recordist and accession numbers are available on DataDryad (<https://datadryad.org/review?doi=doi:10.5061/dryad.f123796>).

**Table S2** Environmental data averaged by study site.

| Site | EVI | VCF | Elevation (m) | BIO1  (°C) | BIO12  (mm) | Distance from contact zone |
| --- | --- | --- | --- | --- | --- | --- |
| Coca, 34 km N of; Napo | 4892.96 | 70.4 | 314 | 24.5 | 3490 | 932 |
| La Selva Lodge,Napo | 5229 | 74.56 | 247 | 25 | 3175 | 887.9 |
| Imuya Cocha; Napo | 5350.88 | 69.16 | 190 | 25.9 | 2680 | 832.5 |
| Tiputini Biodiversity Station | 4790.66 | 74.31 | 232.41 | 25.256 | 3012.81 | 860.2 |
| Loreto, 15 to 80 km W by road; Napo | 3988.56 | 48.96 | 600 | 22.9 | 3674 | 900.6 |
| Canelos; Pastaza | 4539.64 | 75.68 | 507 | 23.4 | 3923 | 821.9 |
| Jau, Parque Nacional de; AM | 4920.64 | 81.16 | 37 | 27 | 2436 | 1017.5 |
| Amani, Lago, AM | 6054.04 | 78.84 | 61 | 26.5 | 3309 | 699.7 |
| El Tigre; Loreto | 5551.2 | 83.52 | 181 | 25.7 | 2735 | 645.2 |
| Kapawi Lodge; Pastaza | 6284.96 | 79.04 | 270 | 25.4 | 2845 | 683.7 |
| Miazal, Morona-Santiago | 5363.44 | 79.4 | 310 | 25.1 | 2593 | 720.4 |
| Libertad, 1.5 km S; Loreto | 5258.96 | 80.6 | 130 | 26.3 | 3056 | 516.5 |
| Jacare Un Bau, AM | 5641 | 77 | 28 | 27.5 | 2298 | 1038.7 |
| Sao Francisco, Fazenda, AM | 4616.52 | 66.88 | 22 | 27.3 | 2409 | 1021.6 |
| Pousada Amazonia, AM | 6047 | 69 | 25 | 27.4 | 2326 | 1029.1 |
| Sucusari, Quebrada; Loreto | 5852.16 | 70.24 | 105 | 26.4 | 2796 | 488.6 |
| San Andres, Loreto | 5508.2 | 58.96 | 101 | 26.4 | 2868 | 448.3 |
| Careiro do Castanho; AM | 5229.12 | 49.92 | 30 | 27.2 | 2170 | 987.6 |
| Tupana | 5125.64 | 83.2 | 28 | 26.9 | 2288 | 924.9 |
| Santa Rita, Loreto | 4678 | 75.12 | 82 | 25.9 | 2708 | 313.1 |
| Palmari, Reserva Natural; AM | 4688.4 | 77.36 | 109 | 25.8 | 2702 | 308 |
| Macaca, Igarape do, AM | 4108.68 | 62.12 | 28 | 26.6 | 2472 | 763 |
| Miazi; Zamora-Chinchipe | 5326.88 | 79.4 | 920 | 22.5 | 2304 | 624.5 |
| Benjamin Constant, AM | 5870.64 | 67.96 | 97 | 25.7 | 2749 | 290.2 |
| Limera, Quebrada; Loreto | 4632.08 | 73.64 | 91 | 26 | 2483 | 340.4 |
| Yarapa Reserve; Loreto | 5327.36 | 77.32 | 107 | 26.5 | 2581 | 362.9 |
| Huampami; AM | 5409.56 | 64.64 | 292 | 25.6 | 2627 | 574.2 |
| Samiria, Rio; Loreto | 5207.64 | 81.8 | 117 | 26.7 | 2381 | 360.1 |
| Angamos, Colonia; Loreto | 4631 | 63.04 | 108 | 26.2 | 2403 | 276 |
| Livramento, Igarape; AM | 4963.4 | 72.36 | 68 | 26.4 | 2379 | 373.5 |
| Labrea, AM | 4926.8 | 65.76 | 62 | 26.4 | 2310 | 366.7 |
| Labrea, 24 km E, AM | 4889.28 | 78.4 | 86 | 26.3 | 2336 | 379.4 |
| Labrea, 36-42 km E, AM | 4921.2 | 79.48 | 93 | 26.2 | 2335 | 357.75 |
| Serra do Divisor, Parque Nacional, AC | 5141.2 | 80.84 | 300 | 25.7 | 2454 | 69.5 |
| Humaita, AM | 4578.36 | 76.36 | 72 | 26.4 | 2206 | 519.3 |
| Abujao; Ucayali | 4819 | 80 | 254 | 25.8 | 1933 | 0 |
| Yarina-Cocha, Lago; Ucayali | 4438.68 | 39.44 | 149 | 26.4 | 1698 | 0 |
| Manoel Urbano to Feijo, AC | 5153 | 75.8 | 220.8 | 25.08 | 2302.8 | 0 |
| Ouro Preto, Igarape, AC | 4709.32 | 80.2 | 297 | 25.5 | 1877 | 0 |
| Manoel Urbano, AC | 4887.5 | 48 | 159 | 25.1 | 2421.5 | 0 |
| Seringal Ocidente, AC | 4838.84 | 57.32 | 243 | 25.7 | 1876 | 0 |
| 35km NE of Tingo Maria; Santa Elena | 4416.28 | 52.52 | 224 | 26.1 | 2545 | 0 |
| Boca do Acre road, AM | 5280 | 68.75 | 191.75 | 25.675 | 1997.25 | 15.7 |
| Tingo Maria, 25 km SW; Huanuco | 4556.24 | 69.28 | 1208 | 20.9 | 1034 | 94.35 |
| Humaita Reserve, AC | 4912.28 | 66.92 | 169.92 | 25.885 | 1945.08 | 0 |
| Abuja, Rio; Pando | 5050 | 73.08 | 117 | 26.4 | 1586 | 198.3 |
| Bujari, AC | 3819 | 54 | 199 | 25.7 | 1948 | 0 |
| Ramal Jarinal, AC | 4049.2 | 30.4 | 210.6 | 25.3 | 1917.2 | 0 |
| Parque Zoobotanico, Rio Branco, AC | 4504.6 | 64.2 | 163 | 26 | 1944 | 0 |
| Parque Chico Mendes, Rio Branco, AC | 4790 | 52.33 | 147.67 | 26.1 | 1932.67 | 0 |
| Catuaba Reserve, AC | 4934.84 | 64.72 | 215.75 | 25.865 | 1921.45 | 0 |
| Puntijao | 5521.92 | 81.76 | 203 | 26.2 | 1778 | 0 |
| Prov. Oxapampa; Distrito Puerto Bermudez | 6346 | 77.6 | 480 | 24.6 | 2606 | 42.9 |
| Cachuela | 3373.84 | 51.75 | 140.33 | 26.6 | 1631.11 | 208.8 |
| San Juan de Nuevo Mundo; Pando | 4517.4 | 36.56 | 163 | 26.3 | 1759 | 113.4 |
| 13km SE of Oventeni; Monte Tabor | 5058.34 | 74.66 | 1134.5 | 22.2 | 2032 | 21.8 |
| Seringal Cachoeira, AC | 4504.25 | 63.5 | 236.25 | 25.55 | 1790 | 0 |
| Ingavi; Pando | 5394.48 | 60.84 | 162 | 26.3 | 1729 | 122.2 |
| Camino Mucden; Pando | 3814 | 39.72 | 262 | 24.9 | 1761 | 0 |
| Inapari Road | 5991 | 51 | 290 | 24.6 | 1665 | 0 |
| Cobija, 12-20 km SW; Pando | 4536.56 | 28.76 | 278 | 24.9 | 1759 | 0 |
| Belen, Madre de Dios | 5200 | 26 | 329 | 24.4 | 1700 | 0 |
| Iberia, | 4632 | 46.5 | 305 | 24.5 | 1730 | 0 |
| Pacahuara Rd | 5013.5 | 43.5 | 297.25 | 24.525 | 1741.75 | 0 |
| Oceania | 4540.33 | 24.67 | 268 | 24.6 | 1747 | 0 |
| Extrema | 5687.57 | 76.57 | 283.43 | 24.714 | 1776.43 | 0 |
| Provencia La Concepcion; Cuzco | 6510.12 | 72.28 | 687 | 25 | 1920 | 0 |
| ARCC, Las Piedras River | 5001.83 | 67 | 248.5 | 25 | 2301.33 | 0 |
| Chive | 5303 | 77 | 201 | 25.6 | 1942 | 17.9 |
| Manu Wildlife Centre; Madre de Dios | 5350.32 | 79.68 | 283 | 25 | 3217 | 0 |
| Cuzco Amazonica Lodge | 5365.28 | 78.44 | 196 | 25.5 | 2130 | 0 |
| CICRA | 5067.03 | 67.81 | 277.44 | 24.825 | 3268.14 | 0 |
| Puerto Maldonado | 2561 | 19 | 190 | 25.5 | 2248 | 0 |
| Heath River Wildlife Center | 5270.5 | 63 | 197 | 25.5 | 2050.5 | 12.2 |
| Posada Amazonas, Tambopata | 5501.5 | 72.17 | 204.75 | 25.367 | 2489.83 | 0 |
| Explorer's Inn; Madre de Dios | 5265.6 | 76.72 | 209 | 25.4 | 2478 | 0 |
| Amazonia Lodge; Madre de Dios | 5192.88 | 70.92 | 591 | 24.4 | 3033 | 23.1 |
| Atalaya and Pilcopata, between; Cuzco | 5386.96 | 71.92 | 533 | 23 | 3125 | 23.7 |
| Patria, Manu Road | 4896.44 | 33.76 | 541 | 24.2 | 3015 | 28.6 |
| Malinowski CP, Tambopata | 5711 | 77 | 196 | 25.3 | 2844 | 0 |
| San Pedro 1 (Cosnipata valley); Cuzco | 3868.2 | 50.6 | 2040 | 18.3 | 1768 | 50.2 |
| Ccolpa de Guacamayos | 5656.76 | 79.56 | 236 | 25 | 3136 | 0 |
| 16km SW of Pilcopata | 4277.88 | 74.8 | 1179 | 21.5 | 3544 | 31.3 |
| Tambopata Research Center | 5904.33 | 78.83 | 236.33 | 25.017 | 3141.33 | 0 |
| San Pedro 2 (Cosnipata valley); Cuzco | 4062.8 | 50.07 | 1799.17 | 19.883 | 2158.83 | 43.8 |
| Curva Alegre; Puno | 4390 | 68.4 | 1442 | 19.8 | 2185 | 0 |
| Chalalan | 4632.8 | 80.56 | 364 | 24.6 | 1890 | 0 |
| Sajta | 5154.72 | 69.36 | 239 | 25.1 | 3070 | 359.1 |

EVI = Enhanced Vegetation Index; VCF = Vegetation Continuous Field (% canopy cover); Bio1 = Annual mean temperature (°C); Bio12 = Annual precipitation (mm).

Data for each specific recording are available on DataDryad (<https://datadryad.org/review?doi=doi:10.5061/dryad.f123796>).

**Table S3** Acoustic measures taken from *Hypocnemis* antbird songs

| Acoustic character (with abbreviation) | | Definition |
| --- | --- | --- |
| 1 | Song duration (D) | Interval between the onset of the first note and the offset of the final note |
| 2 | Note number (NN) | Number of notes in the entire song |
| 3 | Overall song pace (P) | Number of notes in the entire song minus 1, divided by the song duration minus mean note duration (NN-1/D-d_mean_) |
| 4 | Song pace in 1^st^ tercile (P1) | Pace of notes in the first third of the song |
| 5 | Song pace in 2^nd^ tercile (P2) | Pace of notes in the second third of the song |
| 6 | Song pace in 3^rd^ tercile (P3) | Pace of notes in the last third of the song |
| 7 | Pace change 1 (P12) | Change in pace between first and second tercile, calculated as P1/P2 |
| 8 | Pace change 2 (P23) | Change in pace between second and last tercile, calculated as P2/P3 |
| 9 | Mean note duration (d_mean_) | Mean duration of notes, averaged across entire song |
| 10 | Variance in note duration (d_var_) | Variance in note duration across entire song |
| 11 | Mean internote interval (int_mean_) | Mean inter-note interval, averaged across entire song |
| 12 | Internote interval variance (int_var_) | Variance in internote interval across entire song |
| 13 | Mean note maximum frequency (fmax_mean_) | Upper frequency bound of the notes, averaged across the entire song |
| 14 | Mean note minimum frequency (fmin_mean_) | Lower frequency bound of the notes, averaged across the entire song |
| 15 | Mean note peak time (tpeak_mean_) | Time at which peak energy occurs as a fraction of note duration, averaged across entire song |
| 16 | Note peak time variance (tpeak_var_) | Variance in note peak time across entire song |
| 17 | Mean note peak frequency (fpeak_mean_) | Note peak frequency averaged across entire song |
| 18 | Note peak frequency variance (fpeak_var_) | Variance in note peak frequency across entire song |
| 19 | Mean note bandwidth (bw_mean_) | Note bandwidth averaged across entire song |
| 20  21  22 | Note bandwidth variance (bw_var_)  Mean note duration / internote interval (rdint_mean_)  N duration / internote interval variance (rdint_var_) | Variance in note bandwidth across entire song  Mean of ratio of note duration and subsequent internote interval for all notes in the song  Variance of ratio of note duration and subsequent internote interval for all notes in the song |
| Units for temporal characters = seconds; units for spectral characters = hertz (Hz). Measures automatically extracted or calculated from segmented cuts in MatLab. Means of the 22 values were calculated for each individual and used in subsequent analyses. | | |

**Table S4** Loadings of acoustic traits on seven principal components

|  | PC1 | PC2 | PC3 | PC4 | PC5 | PC6 | PC7 |
| --- | --- | --- | --- | --- | --- | --- | --- |
| Eigenvalue | 4.768 | 3.225 | 2.923 | 2.449 | 2.031 | 1.975 | 1.282 |
| % variance | 21.67 | 14.66 | 13.29 | 11.13 | 9.23 | 8.98 | 5.83 |
| D | 0.069 | -0.005 | -0.097 | -0.006 | 0.650 | -0.003 | 0.031 |
| NN | -0.053 | -0.011 | 0.102 | 0.086 | 0.647 | -0.017 | 0.059 |
| P | -0.210 | -0.057 | 0.389 | 0.166 | -0.124 | -0.027 | 0.035 |
| P1 | 0.095 | -0.057 | 0.558 | -0.114 | -0.043 | 0.064 | 0.069 |
| P2 | 0.041 | -0.020 | 0.189 | 0.518 | 0.023 | 0.040 | 0.011 |
| P3 | -0.300 | 0.026 | 0.115 | 0.302 | 0.077 | 0.039 | -0.080 |
| P12 | 0.017 | -0.025 | 0.166 | -0.625 | -0.085 | 0.018 | 0.045 |
| P23 | 0.369 | -0.048 | 0.082 | 0.220 | -0.058 | -0.004 | 0.109 |
| d_mean_ | 0.331 | 0.033 | -0.166 | -0.124 | 0.156 | 0.038 | -0.114 |
| d_var_ | 0.416 | -0.019 | 0.014 | -0.052 | 0.039 | 0.012 | 0.032 |
| int_mean_ | -0.150 | -0.039 | -0.455 | -0.106 | -0.025 | -0.008 | 0.122 |
| int_var_ | 0.157 | -0.063 | -0.424 | 0.324 | -0.230 | 0.077 | 0.072 |
| fmax_mean_ | 0.002 | 0.487 | -0.010 | -0.003 | -0.052 | 0.234 | -0.020 |
| fmin_mean_ | -0.003 | -0.181 | 0.009 | 0.013 | -0.024 | 0.629 | -0.046 |
| tpeak_mean_ | -0.067 | -0.060 | -0.026 | -0.045 | 0.100 | 0.086 | 0.763 |
| tpeak_var_ | 0.169 | 0.114 | 0.066 | 0.078 | -0.138 | -0.255 | 0.537 |
| fpeak_mean_ | 0.018 | 0.194 | 0.017 | 0.016 | -0.005 | 0.592 | 0.083 |
| fpeak_var_ | -0.005 | 0.335 | 0.007 | -0.037 | 0.056 | 0.223 | 0.159 |
| bw_mean_ | 0.014 | 0.539 | -0.011 | -0.018 | -0.035 | -0.052 | -0.006 |
| bw_var_ | -0.027 | 0.507 | 0.001 | 0.046 | 0.043 | -0.239 | -0.096 |
| rdint_mean_ | 0.389 | 0.033 | 0.079 | -0.009 | 0.070 | 0.037 | -0.136 |
| rdint_var_ | 0.443 | -0.003 | 0.087 | 0.077 | -0.041 | -0.024 | -0.051 |

PCA conducted using Varimax rotation. For definition of each abbreviated term, see Table S3. Raw song measurements and PCA scores for each recording are available on DataDryad (<https://datadryad.org/review?doi=doi:10.5061/dryad.f123796>).

**Table S5** Results of ANOVA for each variable compared among subspecies. See table 1 for Tukey HSD results of between subject comparisons.

| Component | Partial SS | df | | MS | | F | *P* | r^2^ |
| --- | --- | --- | --- | --- | --- | --- | --- | --- |
| PC1 | 69.37 | 3 | 23.12 | | 5.02 | | 0.002 | 0.04 |
| PC2 | 147.21 | 3 | 49.07 | | 17.32 | | <0.00001 | 0.13 |
| PC3 | 230.18 | 3 | 76.73 | | 33.47 | | <0.00001 | 0.22 |
| PC4 | 223.5 | 3 | 74.5 | | 40.64 | | <0.00001 | 0.26 |
| PC5 | 45.73 | 3 | 15.24 | | 7.95 | | 0.00004 | 0.06 |
| PC6 | 237.73 | 3 | 79.24 | | 60.3 | | <0.00001 | 0.34 |
| PC7 | 58.49 | 3 | 19.5 | | 17.31 | | <0.00001 | 0.13 |
| PEAK | 0.079 | 3 | 0.026 | | 44.2 | | <0.00001 | 0.27 |
| PACE | 0.23 | 3 | 0.076 | | 44.75 | | <0.00001 | 0.28 |

**Table S6** Generalized linear mixed models (GLMM) of the effect of distance from the contact zone on acoustic structure of the songs of *H. peruviana* (*n* = 198 individuals from 60 sites), with bold denoting significant effects after correcting for false discovery rates [8,9].

| Response variable:  Song structure | Parameter Estimate (**) ± SE | *z* | *P* |
| --- | --- | --- | --- |
| Peak frequency | 0.007 ± 0.023 | 3.06 | **0.002** |
| Overall song pace | 0.068 ± 0.039 | 2.45 | 0.014 |
| PC1 | 0.225 ± 0.177 | 1.27 | 0.205 |
| PC2 | 0.351 ± 0.119 | 2.95 | **0.003** |
| PC3 | 0.456 ± 0.154 | 2.96 | **0.003** |
| PC4 | 0.122 ± 0.095 | 1.28 | 0.201 |
| PC5 | 0.111 ± 0.117 | 0.95 | 0.344 |
| PC6 | 0.272 ± 0.083 | 3.26 | **0.001** |
| PC7 | 0.234 ± 0.055 | 4.27 | **<0.001** |

**Table S7 Geographical and environmental predictors of song structure across the range of *Hypocnemis peruviana*, showing song convergence towards the contact zone**

|  | Acoustic trait | | | | | | | | | | |
| --- | --- | --- | --- | --- | --- | --- | --- | --- | --- | --- | --- |
| Predictors | Peak | Pace | PC1 | | PC2 | PC3 | | PC4 | PC5 | PC6 | PC7 |
| Distance to contact zone | **3.06** | 2.45 | | 1.38 | **2.95** | | **2.96** | 1.27+ | 0.95 | **3.26** | **4.27**** |
| Elevation | 0.62 | 2.28 | | 0.14 | -0.45 | | **3.54*** | 1.33 | -0.31 | 0.10 | 0.47 |
| Annual mean temperature (Bio1) | 0.09 | 1.19+ | | -1.93+ | -0.50 | | 0.79 | 0.87 | -0.75 | 0.02 | **-2.91** |
| Annual precipitation (Bio12) | 1.68+ | 1.33 | | -1.54 | 0.52 | | 0.94 | 0,66 | -0.06 | 2.28 | 0.22 |
| Vegetation structure (EVI) | -1.11 | -0.33 | | 0.32 | 1.26 | | -1.26 | 1.95+ | 1.34 | -2.05 | 0.16 |
| Percent canopy cover (VCF) | -0.56 | 1.16 | | -0.59 | -0.58 | | 1.46 | -0.27 | -1.72 | -0.73 | -0.66 |

Shown are *z*-scores from generalized linear mixed models, with bold denoting significant effects after correcting for false discovery rates using the sharpened method [12, 13]: * *P* <0.01, ***P*<0.001. Models reveal a highly significant effect of distance to the contact zone on a number of key acoustic traits controlling for a range of environmental variables (Table S2). Overall, the songs of *H. peruviana* become more similar to those of sympatric *H. subflava* in spectral traits as they approach the contact zone (see main text). + Denotes variable was included alongside significant factors in best-supported model based on the lowest AIC score. Values are those from the best-supported model, or (when absent) from the original model with all predictors included.
